# Dynamic reconfiguration of macaque brain networks during free-viewing of natural scenes

**DOI:** 10.1101/2021.04.16.439433

**Authors:** Michael Ortiz-Rios, Fabien Balezeau, Marcus Haag, Michael C. Schmid, Marcus Kaiser

**Author notes:** Corresponding author: Michael Ortiz-Rios. Author Contributions: Conceptualization, MOR; Design, MOR; Imaging Methods MOR and FB.; Investigation, MOR, FB & MH; Imaging analysis, MOR; Network fMRI analysis, MOR, MK; Writing – Original Draft, MOR; Writing – Review & Editing, MOR, MCS, MK; Funding Acquisition, MCS; Supervision, MK, MCS.

## Abstract

Natural vision involves the activation of a wide range of higher-level regions processing objects, motion, faces and actions. Here, we pursue a data-driven approach to explore how higher-level visual processes relate to the underlying structural and functional connectivity. Using a free-viewing paradigm in four awake rhesus macaque monkeys, we investigate how different visual scenes change functional connectivity. Additionally, we explore how such functional connectivity, as measured through fMRI, is related to the structural connectivity, as measured through diffusion weighted imaging. At first, we evaluate the consistency of the elicited free-viewing pattern using standard analytical techniques. We also evaluate the underlying structural connectivity via diffusion data by tracking white matter bundle projections from the visual cortex. We then reconstruct free-viewing and structural networks and quantify their properties. Centrality measures over the entire fMRI time-series revealed a consistent functional network engaged during free-viewing that included widespread hub regions across frontal (FEF, 46v), parietal (LIP, Tpt), and occipitotemporal cortex (MT, V4 and TE) among others. Interestingly, a small number of highly-weighted and long-length inter-hemispheric connections indicated the presence of long-range integrative properties during free-viewing. We hypothesized that during free-viewing, networks had the capacity to change their local and distal connections depending on the on-going changes in visual scenes. To capture these network dynamics, we depart from the static modular architecture of the structural networks and demonstrate that hubs in free-viewing networks reorganize according to the presence of objects, motion, and faces in the movie scenes indicating poly-functional properties. Lastly, we compare each NHP subject network and observe high consistency between individuals across the same network type with closer correspondence between structural networks (e.g., diffusion based and those partially assembled from tract-tracing). In summary, our network analyses revealed ongoing changes in large-scale functional organization present during free-viewing in the macaque monkey and highlight the advantages of multi-contrast imaging in awake monkeys for investigating dynamical processes in visual cognition. To further promote the use of naturalistic free-viewing paradigms and increase the development of macaque neuroimaging resources, we share our datasets in the PRIME-DE consortium.

## 1 Introduction

Recent progress in multimodal neuroimaging has enabled a new paradigm shift that emphasizes connectivity as the core of brain organization and places inter-regional communication as a central framework for understating cognition. Previous connectivity studies in humans established that brain signals transit between states of high and low connectivity strength over time and that these fluctuations relate to patterns of synchrony in the network (Zalesky et al., 2014). But how these state fluctuations relate to internal or external triggers for reconfiguration remains unknown and challenging to infer during resting state paradigms.

More recently, an increasing number of studies, in both human and monkeys, began to implement naturalistic viewing paradigms for mapping large-scale changes in brain activity with rich temporal dynamics (Bartels and Zeki, 2004a, 2005; Bartels et al., 2008; Hung et al., 2015; McMahon et al., 2015; Russ and Leopold, 2015; Russ et al., 2016). For example, the recent macaque studies by Russ and Leopold, 2015 and Sliwa and Freiwald, 2017a demonstrated how the free-viewing paradigm could be used for mapping specialized areas without the use of trial-by-trial experimental designs.

In humans, previous studies using the naturalistic free-viewing imaging paradigm began to establish the degree of similarity between activation maps obtained during movie viewings and maps obtained with more controlled stimulation (Bartels and Zeki, 2004b, 2005; Bartels et al., 2008). Further fMRI studies demonstrated how functional connectivity patterns were synchronized across individuals watching the same movie clips (Hasson et al., 2004a; Lahnakoski et al., 2012), highlighting the reliability and effectiveness of the approach. Additional studies compared the activation patterns between humans and macaque monkeys watching identical movies and identified homologous and divergent brain regions between primate species (Betti et al., 2013; Mantini et al., 2012). More detailed mapping of the movie viewing activation patterns in monkeys also revealed a previously unknown predominance of visual motion-related activation in areas well-described for face processing (Russ and Leopold, 2015; Russ et al., 2016) as similarly found in humans (Hasson et al., 2004b). Interestingly, recent graph-theoretical network studies in humans have revealed that face-networks tend to reorganize when inverted faces are viewed (Rosenthal et al., 2017), reinforcing the notion that inferotemporal face networks might be more dynamic in their representational state than previously conveyed using trial-by-trial designs. Importantly, in humans the free-viewing approach applied in combination network analytical tools is beginning to highlight important insights of brain function that could not be easily revealed using standard trial-by-trial or resting-state paradigms (Betzel et al., 2016).

In the macaque, most network studies have focused on the compilation of anatomical tract-tracing datasets for inferring signal propagation from the underlying structural “backbone” of the network (Harriger et al., 2012; Markov et al., 2014). Moreover, most functional connectivity studies in NHPs have practically relied on the use of resting-state paradigms largely carried out under general anesthesia (Hayashi et al., 2021). Here, we implement a free-viewing approach for describing the relationships across brain regions in four macaque monkeys. Importantly, we avoid the use of general anesthesia and contrast enhancement agents, but relied instead on the classical blood oxygen level depended (BOLD) signal as commonly done in functional connectivity studies in humans (Hutchison et al., 2013). Given the challenges of acquiring awake macaque neuroimaging datasets, our work here began as an exploratory inspection of the quality of the datasets. Moreover, since free-viewing networks have not yet been characterized, we consider our study here as important first step in establishing the acquisition and analytical methods for investigating the dynamical nature present in macaque brain networks.

Towards this end, we began exploring our data using multiple analytical approaches (e.g., general linear modeling (GLMs), independent component analysis (ICA) and coherence analyses) as a step to validate functional imaging data. For diffusion weighted imaging (DWI) data, we first validated the quality of our tractography by tracking white matter bundles pathways from visual cortex. After these initial analyses, we then reconstruct structural networks based on the number of streamlines touching every pair of ROIs in the network (Bassett and Sporns, 2017; Bullmore and Sporns, 2009), while free-viewing networks were constructed from correlations across the regional mean BOLD signals. We then characterized free-viewing networks and structural networks by measuring node degree, node degree distribution, path length, path length distribution, clustering coefficient, centrality and modularity. Importantly, from structural networks we classify regions based on their static modular architecture which enables the inference of network changes during free-viewing. Additionally, we compare networks across modalities (e.g. free-viewing networks versus structural networks) and further compare networks from our four macaques with those assembled from openly available tract-tracing data (Markov et al., 2014). From a second movie experiment, we investigated the extent to which free-viewing networks reconfigure by applying time-varying functional connectivity to the rich time-series and observed specific cortico-cortical and thalamocortical functional interactions during each main epoch. We discuss our findings in relation to the well-known functional and anatomical properties of the macaque visual system and discuss how natural-viewing paradigms in NHPs in combination with network neuroscience tools can help us understand the dynamical nature of large-scale cognitive systems. We believe that our datasets could be beneficial to the neuroimaging community for allowing cross-species comparisons (e.g., human and monkey) of free-viewing states. To this end, we share our raw, pre-processed and free-viewing networks datasets following the guidelines of the PRIME-DE consortium, to further promote the development of new complementary neuroimaging resources to resting-state.

## 2 Materials and Methods

### 2.1 Subjects

Four female rhesus monkeys (*Macaca mulatta*) were used to obtain all neuroimaging data; (VL, six years of age, weighing 7 kg; DP, six years of age, weighing 9 kg; FL 4 years of age, weighting 6 kg; AL, five years of age, weighing 9 kg). All animals were socially housed and were environmentally enriched in their home cage.

A very important facet of NHP neuroimaging, as opposed to human neuroimaging, is the requirement for head stabilization during data acquisition. Head stabilization in NHPs is accomplished via head post implants. The size and material of such implants affect the quality of functional images near the implantation region which further affects the reliability and construction of whole-brain networks from imaging data. Here, we applied a customized head-post implant procedure on all monkeys that allowed us to obtain distortion-free MR-images across the whole brain. All surgical and anesthesia procedures, postoperative care, and implant methods were described in detail previously (Ortiz-Rios et al., 2018b).

The UK Home Office approved all procedures, complying with the Animal Scientific Procedures Act (1986) for the care and use of animals in research and the European Directive on the protection of animals used in research (2010/63/EU).

### 2.2 Awake macaque neuroimaging

Prior to data acquisition, monkeys were trained to voluntarily approach the inside of a wooden box from which they then learned to enter a cylindrical MRI-compatible chair. Once inside the chair, the animals were trained to move their head outside of the chair and to remain calm with their head outside of the chair by using positive reinforcement techniques.

Once habituated, animals were then exposed to the scanner environment and were acclimatized to remain calm. Acclimatization within the scanner took one month to accomplish. Once animals were head implanted and were comfortable with the procedure, the animal head was immobilized in the MRI chair using (two months post implantation) with a PEEK head holding system apparatus. The monkey’s ears were protected from the scanner noise by using standard ear muffs. During the scanning period, animals remained calm while watching five minutes movie clips. Typically, while being trained or scanned in the absence of any visual stimulation, in darkness or during anatomical scanning animals remained calm or on a sleep calm condition. Acclimatizing the macaques for awake imaging took approximately three months to accomplish.

#### 2.2.1 Stimuli presentation

A 45-degree angled back-projection mirror system displayed all movie stimuli. A projector (NEC NP1150) projected images onto a screen (size: 35 length x 32 width cm) which was placed at a distance of 73.5 cm from the center of the monkey’s head. The projection created a visual field of 26 x 24 degrees of visual angle.

An infrared based eye-tracking system (iView, SensoMotoric Instruments GmbH, Teltow, Germany) received a video signals from an MRI-compatible camera (12M-I with integrated LED, MRC Systems, GmbH) placed behind a mirror in front of the NHPs eye. An analog-to-digital conversion card (NI-USB 6212-BNC, National Instruments) then sampled the analog signal from the eye-tracker. Additionally, the digital-to-analog output of the NI-card was used to control the reward system and used to trigger the start of data acquisition. The NI-card was controlled via MWorks software (<u>the MWorks project</u>) running on a MacMini (2.8 GHz, Core i5).

#### 2.2.2 Movie experiment one

The initial movie clips contained unfamiliar visual scenes with ego-perspective camera motion. The movie clips showed an ego perspective scene of a human individual driving a bike across a rough terrain (see https://github.com/ortizriosm/natural-vision-connectivity.). The next two segments showed two human individuals climbing a transmission power tower. Within an imaging run, five movie clips (30 secs on and OFF) were presented, lasting a total of five minutes. During the OFF period the screen turn dark. During the movie presentation monkeys were not rewarded and received no liquid to avoid jaw movements during imaging. Animals were rewarded using juice reward after the movie ended and once the scanner acquisition period had finished. The content of the scenes were kept constant across repetitions and sessions for all four macaques allowing us to compare activation patterns across all four subjects.

#### 2.2.3 Movie experiment two

For the second series of free-viewing experiments, each movie segment lasted 30 seconds, followed by a dark interval period of 15 seconds. We presented twenty-five movie segments (see https://github.com/ortizriosm/natural-vision-connectivity.) in a run which lasted ∼20 minutes. Each movie type was modified to control the low-level visual features on a frame-by-frame basis using the following methods. For the phase scrambling the original frames were Fourier transformed using Matlab, then the phase was randomized and added to the initial phase. The inverse Fourier was then calculated over the transformed data to generate an image. For the other three controls, we used the Matlab functions *randblock* for tile scrambling, the *spectral visual saliency toolbox* to create saliency contour images and *OpticalFlow Matlab* function to create vector motion direction on each frame.

### 2.3 Multimodal data acquisition

A vertical 4.7 Tesla magnet running ParaVision 5.1 (Bruker, BioSpin GmbH, Ettlingen, Germany) and equipped with a 4-channel phase-array coil covering the whole head (https://www.wkscientific.com) was used to acquire MR images. Imaging data consisted of three types of datasets: Diffusion weighted imaging (DWI) for white matter bundle tractography, echo-planar imaging (EPI) for functional analyses and T1 for anatomical segmentation and parcellation (**Table 1** and **Fig.1A**).

**Figure. 1.**
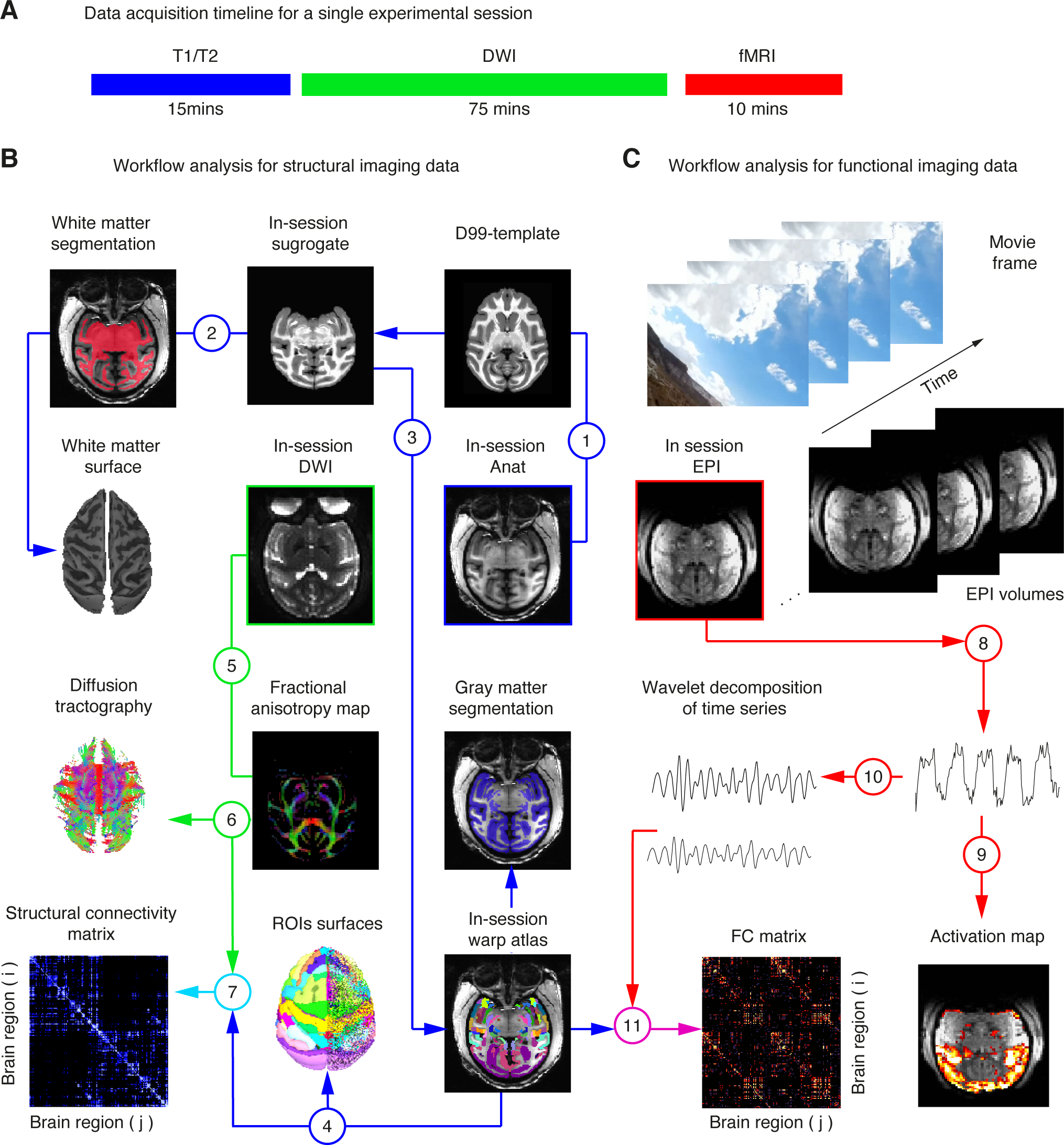
Flow chart for the acquisition and analysis of structural and free-viewing networks. **A**. Workflow for the acquisition of structural T1, T2, DWI and fMRI datasets. All data was approximately acquired within less than two hours of scanning time. **B**. Data pre-processing flow chart for the construction of Structural networks. **C**. Data pre-processing flow chart for the construction of Free-viewing networks. We acquired three imaging datasets within a single experimental session: Diffusion-weighted imaging data (DWI, green), anatomical (T1, blue), and the echo-planar imaging data (EPI, red). The color-coded arrows show the pre-processing steps for each type of dataset as follows: (*1) Warping the anatomical data into the D99 template to generate an in-session subject surrogate brain; (2) White matter segmentation and surface reconstruction; (3) Atlas parcellation aligned to the In-session anatomy and grey matter segmentation; (4) Rendering of ROIs; (5) DWI standard pre-processing; (6) Diffusion tractography and rendering of streamline tracts; (7) Structural networks construction based on the pairwise number of streamline connections; (8) Standard fMRI pre-processing; (9) GLM and coherence analyses; (10) Wavelet decomposition (frequencies 0.04 - 0.16 Hz); (11) Free-viewing networks construction from pairwise cross-correlation of the mean time series of each ROI.* See **Methods** section for a detailed description of each step.

**Table 1.** Acquisition parameters for each subject session and of type dataset: DWI, EPI and MDEFT. (Left) a diffusion-weighted spin-echo EPI sequence (DW SE-EPI) sequence was used to acquire DWI images for white matter tractography. (Middle) a gradient-echo (GE) EPI sequence was used to acquire BOLD signal modulation for functional imaging. (Right) a magnetization-prepared RApid gradient echo (MP-RAGE) sequence was used to acquire anatomical (T1) images for cortical white and gray matter segmentation and atlas parcellation. Nacq, (Number of acquisitions); Acl. Fac., (Acceleration factor); Seq. (Sequence), Bo im., (number of B0 images); B dir. (number of B directions).

| Subj.Sess. | DWI |  |  |  | EPI |  |  |  | MDEFT |  |  |  |
| --- | --- | --- | --- | --- | --- | --- | --- | --- | --- | --- | --- | --- |
|  | V.Nr1 | D.lp1 | F.O41 | A.O01 | V.GL1 | D.GI2 | F.MB1 | A.Pr1 | V.GL1 | D.GI2 | F.MB1 | A.Pr1 |
| FA (°) | 65 | 65 | 65 | 65 | 65 | 65 | 65 | 65 | 90 | 90 | 90 | 90 |
| TE (ms) | 58 | 58 | 58 | 58 | 21 | 21 | 21 | 21 | 3.74 | 3.74 | 3.74 | 3.74 |
| TR (ms) | 14200 | 14200 | 14200 | 14200 | 1500 | 1500 | 1500 | 1500 | 2000 | 2000 | 2000 | 2000 |
| BW<br>(Hz/Px) | 1495 | 1495 | 1495 | 1495 | 1704 | 1704 | 1704 | 1704 | 284 | 284 | 284 | 284 |
| ES (ms) | 58 | 58 | 58 | 58 | 58 | 58 | 58 | 58 |  |  |  |  |
| Acl. Fac. | 2 | 2 | 2 | 2 | 2 | 2 | 2 | 2 | 1 | 1 | 1 | 1 |
| Seq. | Dw-SE-EPI | Dw-SE-EPI | Dw-SE-EPI | Dw-SE-EPI | GE-EPI | GE-EPI | GE-EPI | GE-EPI | MDEFT | MDEFT | MDEFT | MDEFT |
| B val<br>(s/mm2) | 800 | 800 | 800 | 800 | x | x | x | x | x | x | x | x |
| Bo im. | 4 | 4 | 4 | 4 | x | x | x | x | x | x | x | x |
| B dir. | 60 | 60 | 60 | 60 | x | x | x | x | x | x | x | x |
| RR-RO<br>(mm) | 0.97 | 0.97 | 0.97 | 0.97 | 1.2 | 1.2 | 1.2 | 1.2 | 0.61 | 0.61 | 0.61 | 0.61 |
| RR-PH<br>(mm) | 0.97 | 0.97 | 0.97 | 0.97 | 1.2 | 1.2 | 1.2 | 1.2 | 0.61 | 0.61 | 0.61 | 0.61 |
| RR-SL<br>(mm) | 1 | 1 | 1 | 1 | 1.2 | 1.2 | 1.2 | 1.2 | 0.62 | 0.62 | 0.62 | 0.62 |
| RIM-RO<br>(px) | 88 | 88 | 88 | 88 | 88 | 88 | 88 | 88 | 176 | 176 | 176 | 176 |
| RIM-PH<br>(px) | 88 | 88 | 88 | 88 | 88 | 88 | 88 | 88 | 176 | 176 | 176 | 176 |
| RIM-SL<br>(px) | 56 | 56 | 56 | 56 | 31 | 34 | 34 | 34 | 72 | 74 | 96 | 74 |
| Nacq | 5 | 5 | 5 | 5 | 200 | 200 | 200 | 200 | 1 | 1 | 1 | 1 |
| TA<br>(h:m:s) | 01:15 | 01:15 | 01:15 | 01:15 | 05:00 | 05:00 | 05:00 | 04:24 | 14:24 | 14:48 | 19:12 | 04:24 |

### 2.4 Overall pipeline workflow for the analyses of multimodal data

To provide clarity, we illustrate the workflow of data pre-processing and analyses for the construction of networks in **Fig 1A** and **1B**. The analyses consisted of two pre-processing steps for each dataset type: T1 (blue in steps 1 – 4) DWI (green 5 – 7) and EPI (red in steps 8 – 10). The construction of structural networks (magenta in step 7) and the construction of free-viewing networks (pink in step 11) were the final result of the pre-processing pipeline. All pre-processing analyses were conducted using openly software provided in the packages of AFNI (Cox, 1996a), SUMA (Saad et al., 2004) and Freesurfer (Dale et al., 1999). Additional analyses for functional connectivity and network measures were performed using the functions provided by the Brain Connectivity Toolbox (BCT) https://sites.google.com/site/bctnet/Home and Matlab.

### 2.5 Anatomical data pre-processing and atlas parcellation

For the in-session anatomy (T1), the D99 (Reveley et al., 2017) atlas was warped into the in-session T1 volume, generating an in-session surrogate brain. We used the *align_macaque_script* (Step 1) to generate all atlas files from which then we optimize the integration of some of the very small ROIs for network construction. For the white matter segmentation (Step 2), the command *3dSeg -anat “input.nii” -mask AUTO -classes output.nii -bias_classes “output.ni” - bias_fwhm 25 -mixfrac UNI -main_N 5 -blur_meth BFT,* was used to obtain a binary white matter (WM) mask. The output from *align_macaque_script* also included the atlas ROIs parcellation of the D99 template (Step 3). The atlas file was then used for segmentation of the cortical grey matter by binarizing the atlas file using the programs *3dAutomask* and *3dcalc* (Step 4). The masks were then used to generate a rendered white, gray (Step 5) and inflated surfaces (Step 6) in FreeSurfer using the autorecon-wm commands. For displaying the ROIs into surface, we first created a color mappable niml file of the atlas volume using the command, *3dVol2Surf -spec input.spec -surf_A lh.surface.gii -sv input.nii -grid_parent rois.nii - use_norms -norm_len 2.5 -map_func max -f_steps 10 -f_index nodes -out_niml Lh.output.niml.dset*.

We then display the activation results along with ROIs using contours.

### 2.6 fMRI data pre-processing

All fMRI data was pre-process using (Analyses of functional neuroimages – AFNI) (Cox, 1996b). Pre-processing of fMRI time series (Step 8) included: slice-timing correction, motion correction, spatial-smoothing and normalization (a voxel-by-voxel scaling of the time series by the mean). FMRI time-series were corrected for slice-timing differences using the command *3dTshift -tzero 0 -prefix “output.nii” “input.nii”*.

For the occurrence of spikes – transient artefacts often caused by small electrical discharges or monkey movement – we detected them using the command, *3dDespike -NEW25 -localedit -prefix “output.nii” “input.nii”*.

For dispiking a linear regression curve is fitted into the time-series and the mean absolute deviation (MAD) of the residuals is calculated. The fraction of outliers was then calculated from the time-series using the command, *3dToutcount -automask “input.nii” > “output.1D”*.

Next, we performed motion correction by first calculating a mean baseline volume suing the command, *3dTstat -prefix “epi.mean.nii” “input.nii”*.

Motion correction was then performed with the command, *3dvolreg -verbose -base “epi.mean.nii” -dfile “output.1D” -prefix “output.nii” “input.nii”*.

*3dvolreg* applied a rigid body transformation with 6 parameters (3 translation and 3 rotation) to each time-point in the time-series to match the voxelwise mean obtained over the entire time-series in each volume. The output of (*3dvorleg*) contained motion parameters (output.1D file) which were input into regression analyses as nuances of no-interest. Using the python script, *1d_tool.py -infile input.1D -set_nruns 1 -show_censor_count -overwrite -censor_motion 0.5 “output.1D”*, we detected motion shifts greater than 0.5 mm and rotations greater than 0.5 degrees. More than 98% of the volumes were included in the analyzes since movement deviations were minimal or non-detectable from movement derivatives of the time-series. Derivatives of the motion parameters were calculated using the command, 1d_tool.py -infile “input.1D” -set_run_lengths 200 -set_tr 1.5 -derivative -overwrite -write “output.1D”.

To improve signal-to-noise ratio and reduce anatomical differences we performed spatially smoothing by convoluting a Gaussian kernel filter with a size of 2.4 mm full-width-half-max (FWHM). The command, *3dmerge -1blur_fwhm 2.4 -doall -prefix “output” “input”,* was used for spatial smoothing and the output data was used mainly for in-volume visualization. In order to select brain voxels from the time-series a brain mask was created using command, *3dAutomask -peels 5 -prefix “output.nii” “input.nii”*.

The brain mask enabled the removal of the surrounding non-brain tissue. Next, we calculated a mean epi from the skull-stripping datasets for each run and calculated a grand average mean epi for all the in-session data. We then used the mean epi mean to calculate a non-linear warp between the mean epi data and the in-session anatomy using the command, *3dQwarp -source “input.anat.nii” -base “input mean.epi.nii” -prefix “output.nii” -mi -verb - iwarp mi -blur 0 3 -Qfinal -verb -blur 0*.

The inverse transformation warp was then applied to the original time series using the command, *3dNwarpApply -nwarp “WARPINV.nii” -source “input.nii” -master “input.nii” -prefix “output.nii”*.

After non-linear alignment, all data was then ready for the first pass regression using analyses with 3dDeconvolve.

### 2.7 General linear-modeling (GLM) analyses

For the first series of experiments, we presented two movie segments that were partitioned into five movie clips each lasting 30 seconds and each followed by a 30 second period of no visual stimulation (darkness). Five movie segments (30 sec each) were presented within a run for a total of 5 minutes per scan (300 sec, 1.5 sec TR, n volumes = 200). As the input dataset to (3dDeconvolve Step 9), we provided the detrended time-series and then applied ordinary least squares to a regression model of the BOLD signal which included a gamma variate block design of the hemodynamic response to the movie times presentation. A general linear T-tests were performed between movie periods and the baseline/blank periods for movie clips. Stimulus times were created with the script (*make_stim_times.py*) which included a binary file containing the respective number of time points for movie presentation and baseline (e.g., 30 secs ON/ TR = 1.5 sec = 20 volumes ones and 20 volumes zeros). The overall block design for movie presentation was initially visualized with the commands (*3dDeconvolve* and *1dplot)* and then submitted for general linear modelling. The stats output from 3dDeconvolve was then used to visualize the activation results. The threshold was chosen to be at a significant and corrected false discovery rate (FDR q value < 0.05). We confirmed significant activation (T-value colormap range 2.3 < 10, FDR q < 0.05) across the visual and higher-visual-related regions in all four macaque monkeys (**Supp. Fig.1A**). These analyses were performed during data acquisition as a strategy to evaluate the ability to get a significant BOLD response and decide whether or not more was required. Overall, we observed a classical general pattern of activation in all four macaque monkeys and in regions previously reported on previous NHPs fMRI experiments (Goense et al., 2012; Logothetis et al., 1999).

### 2.8 Diffusion data pre-processing

For the DWI data, the volumes were first corrected for slice-timing differences using the command, *3dTshift -tzero 0 -prefix input.nii*.

We then averaged the five-volume repetitions on each dataset using the command, *3dcalc -a input.nii’[0..63]’ -b input.nii’[64..127]’ -c input.nii’[128..191]’ -d input.nii’[192..255]’ input.nii’[256..319]’ -expr ’(a+b+c+d+e)/5’ -prefix “output.nii”*.

The average dataset was then aligned to a reference image using the command, *3dAllineate -base “input.nii[0]’ -input input.nii -prefix output.nii -cost mi -verb -EPI*. We then selected the 64 directional data using the command, *3dcalc -a input.nii’ [3..63]’ -expr ’a’ -prefix output.nii,* and entered the output into 3dDWItoDT (Step 5) to compute the six direction vectors (Dxx, Dxy, Dyy, Dxz, Dyz, and Dzz) and reference gradient vectors (Gxi, Gyi, and Gzi). We then executed the command, *3dDWItoDT -prefix output.nii -automask -reweight -verbose 100 -sep_dsets -eigs bvecs input.nii,* to get all the vectors. Additionally, 3dDWItoDT also provided eigenvalues (L1, L2, and L3), eigenvectors (V1, V2, and V3), fractional anisotropy (FA), mean diffusivity (MD), and radial diffusivity (RD). The FA map was transformed into an RGB color-coded map using the command, *3dThreetoRGB -prefix output.nii -anat input.nii’[0]’*.

The results showed the color-coded FA and overall white matter fiber bundles of each monkey (each monkey **Supp. Fig.1B**). We then computed the uncertainty via jackknife of DTI estimates on-voxel-by-voxel basis using the command, *3dDWUncert -inset input.nii -mask input.nii -prefix output.nii -input input.tensors -grads bvecs -iters 50*.

To render the streamline tracts from the visual cortex were then tracked white matter bundle pathways using the program 3dTrackID (Step 6). 3dTrackID is based on fiber assessment by continuous tracking including diagonals (Taylor and Saad, 2013a). For diffusion tractography we implemented the command, *3dTrackID -mode MINIP -dti_in DTI.tensors -dti_extra mask.input.nii -netrois input.rois.nii -logic AND -mini_num 5 -uncert input.nii -alg_Thresh_FA 0.45 -prefix output.nii* Specifically, for tracking bundles in visual cortex, a pair of ROIS were selected from the same hemisphere in atlas via their numerical index. For tracking the optic radiation, we used the LGN as a source and V1 as a target ROI. For tracking the forceps major both V1 ROIS were selected for source and target projection tracking.

### 2.9 Independent component analysis (ICA)

Run-level ICA was performed using the Multivariate Exploratory Linear Optimized Decomposition into Independent Components (MELODIC: http://fsl.fmrib.ox.ac.uk/fsl/fslwiki/MELODIC) ICA module of the FSL package. MELODIC was used to obtain general patterns of BOLD activity that largely project free-viewing networks . ICA estimates the consistency of a set of spatially and temporally overlapping components over the fMRI time-series. Components might consist of meaningful organizing patterns such as those typically measured during resting state conditions in addition to other artifactual effects resulting from head motion, heart pulsation or respiration and each carrying an independent spatial pattern and time course. For our naturalistic paradigm we aimed to obtain the overall meaningful pattern observed during free-viewing form the spatiotemporally rich signals. The MELODIC ICA algorithm attempts to segregate the spatial overlap between the components based on the independence of the fMRI-BOLD signals. ICA is a “model-free” algorithm that aims to detect cortical and subcortical responses that are prevalent among a cluster of voxels, instead of using a modeled BOLD response for comparing the fMRI signal (Hyvärinen and Oja 2000). Previous NHPs studies suggested that the optimal number of components lies within the range of 20–30 independent components for RS-fMRI data (Hutchison et al. 2011; Belcher et al. 2013; Mantini et al. 2013). Here we mainly aimed at replicating some of our previous results with GLM by using ICA. All runs across monkeys showed the predominance of the free-viewing paradigms block design as the main first component detected which explain 7 – 14 % of the variance (see **Supp. Fig.2** and **Table 2**).

**Figure 2.**
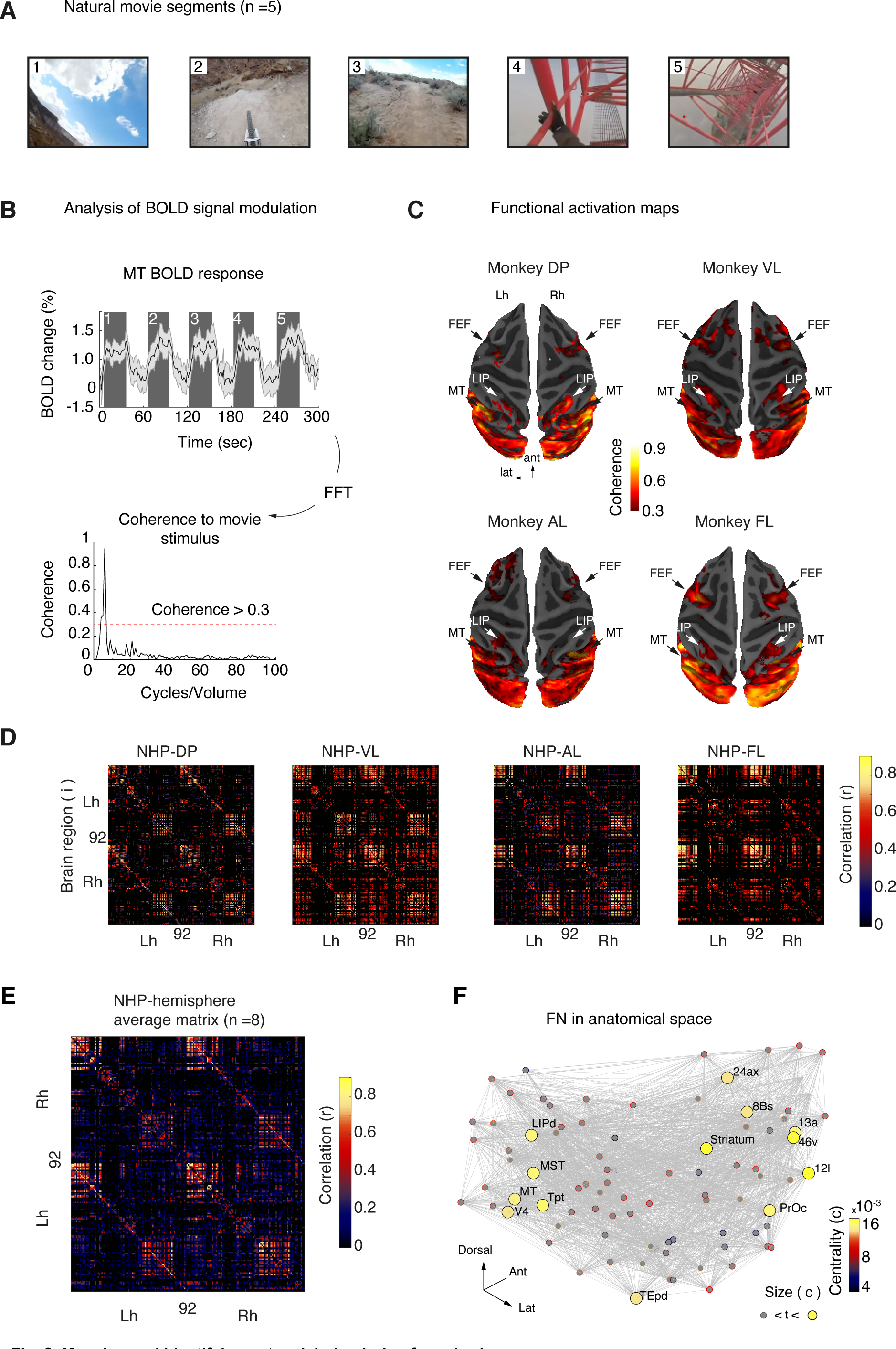
Mapping and identifying network hubs during free-viewing. **A**. Block design paradigm with example frames of each movie segment. Movie segments lasted for 30 seconds, followed by 30 seconds of darkness. **B**. Time course (mean ± SEM) of MT voxels from NHP DP showing BOLD responses to each movie segment. Gray shading represents dark periods within an imaging run. The bottom panel shows the Fourier transform of a voxel showing the BOLD response peak at the stimulation rate (0.016 Hz = 1 cycle/60 s). To calculate coherence, we used the voxel peak response and the stimulation rate. **C**. Resulting coherence maps rendered into a semi-inflated brain surface of each subject. Across all NHPs, regions with coherence modulation (> 0.35) included regions in frontal, parietal, visual, and higher-level visual regions along the inferotemporal gyrus, among others. **D**. Individual free-viewing networks matrices show the absolute correlation coefficient between every pair of mean time series of each ROI. **E**. Average free-viewing network (92 x 92) matrix taken across subjects and hemispheres (n = 8). **F**. Hubs with high eigenvector centrality highlight regions with high-importance in the visual network during natural free-viewing. Identified hub regions highlight a visuo-saccadic network.

### 2.10 Temporal signal-to-noise ratio

To further demonstrate the quality of the awake macaque EPI data (**Supp. Fig. 2**), we calculated the temporal signal-to-noise ratio (tSNR). The tSNR was defined by the mean signal divided by the standard deviation of the signal over the voxel time series (Welvaert and Rosseel, 2013). We used command, *3dTstat -tsnr -prefix output.nii input.nii* to calculate the tSNR on the minimally preprocessed time series (e.g., after alignment to the anatomy). The temporal tSNR maps range between 0 and clips at 100 for easy visual comparison across subjects (**Supp. Fig.3**). Additionally, we calculated the mean TSNR for each run acquisition which ranged between 46 – 82 across subjects and runs (see **Table 2** for details). In general, the maps show high tSNR across gray matter structures with more increases observed in temporal regions that were close to the phase array coil loops.

**Figure. 3.**
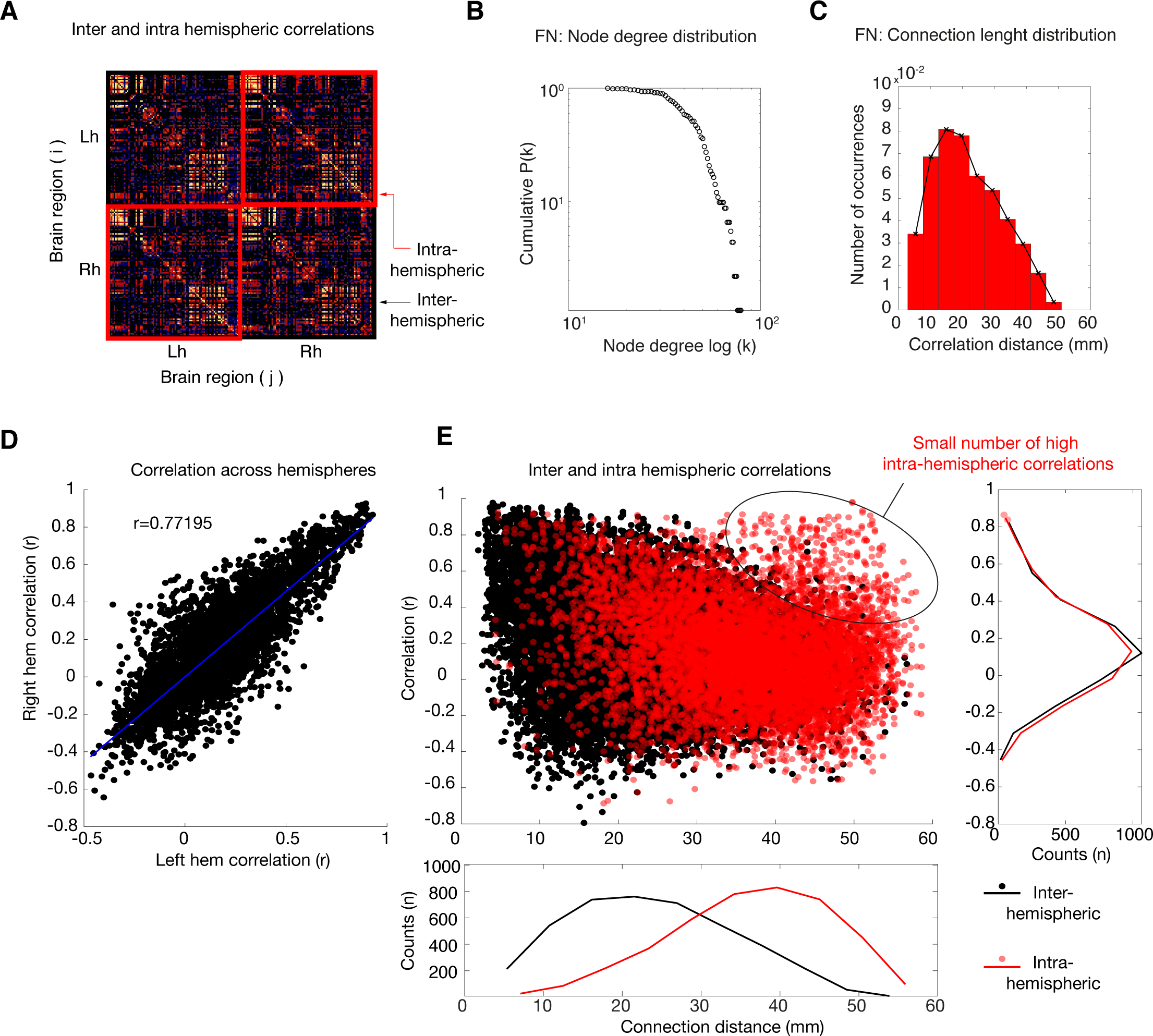
Functional organizing properties of free-viewing networks Organizing properties of Free-viewing networks. **A**. Connectivity matrix organized from left to right hemispheres. **B**. The complementary cumulative distribution function (cCDF) of node degree shows a decrease in the probability of finding a highly connected node (*k*). **C**. Distribution of path lengths shows a decay as a function of Euclidean distance. **D**. Scatter plot of left and right hemisphere correlations showing high similarity across hemispheres. **E**. Scatter plot of node distance and correlation magnitude showing inter (black) and intra (red) hemispheres correlations. The outset panel to the right shows the average counts of correlation coefficients. The outset bottom panel shows the average counts of distance lengths. Both correlations decrease as a function of distance. However, a small number of long-distance inter hemispheric correlations remains.

### 2.11 Coherence analyses

Coherence measures the ratio of the amplitude at the fundamental stimulation frequency to the signal variance, ranging between 0 and 1 (Brewer et al., 2002). The measure of coherence is

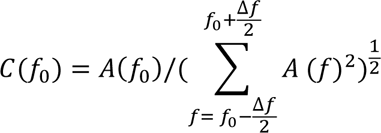

where *f*_0_ is the stimulus frequency, *A*(*f*_0_) the amplitude of the signal at that frequency, *A*(*f*) the amplitude of the harmonic term at the voxel temporal frequency *f* and Δ*f* the bandwidth of frequencies in cycles/scan around the fundamental frequency *f*_0_. For all movie stimuli, *f*_0_ corresponds to one cycle (1/60 sec = 0.016 Hz) and Δ*f* corresponds to the frequencies around the fundamental (see **Fig. 2B**). The threshold was chosen at a coherence level > 0.35. For all monkeys, the movie scenes within a single 5-minute-long scan (2 scan repetitions) elicited a significant BOLD response modulation (coherence > 0.35) in a large number of cortical areas, including occipital, temporal, parietal and frontal regions (**Fig. 2C**).

### 2.12 Construction of free-viewing networks

ROI time series were obtained using the program (3dNetCorr). The anatomical based ROIs were down-sampled to match the underlying EPI grid (3dFractionize) from which each masked ROI was used to obtain the mean time series. The time series output from 3dNetCorr was then read into Matlab for connectivity measures.

Previous studies in humans have utilized maximal overlap discrete wavelet transform (MODWT) for connectivity studies of resting-state fMRI and task-based conditions. Similarly, we used the MODWT to decompose each mean regional time series into wavelet scales corresponding to specific frequency bands (Sizemore and Bassett, 2018). The time-series were wavelet decomposed using the orthogonal *Daubechies* wavelet, which resulted in five level decompositions ranging from 0.001 to 0.25 Hz. For the construction of the functional matrices, we concentrated on two levels (0.125 – 0.06) and (0.06 – 0.03) since the hemodynamic events occurring within the movie viewing periods were within the relatively low-frequency range of 6 to 24 seconds. The function is available at file exchange website (https://uk.mathworks.com/help/wavelet/ref/modwt.html). To compare Free-viewing networks across different wavelet levels we plotted each average matrix for each decomposition (**Supp. Fig 4**). Overall, the FVN pattern on the matrices was presented for much of the level decomposition while the noise level increased for higher frequency levels and decreased for lower frequency levels. The correlation coefficients between the original matrix and levels 2 and 3 were highest, while lowest for the high-frequency level 5.

**Figure. 4.**
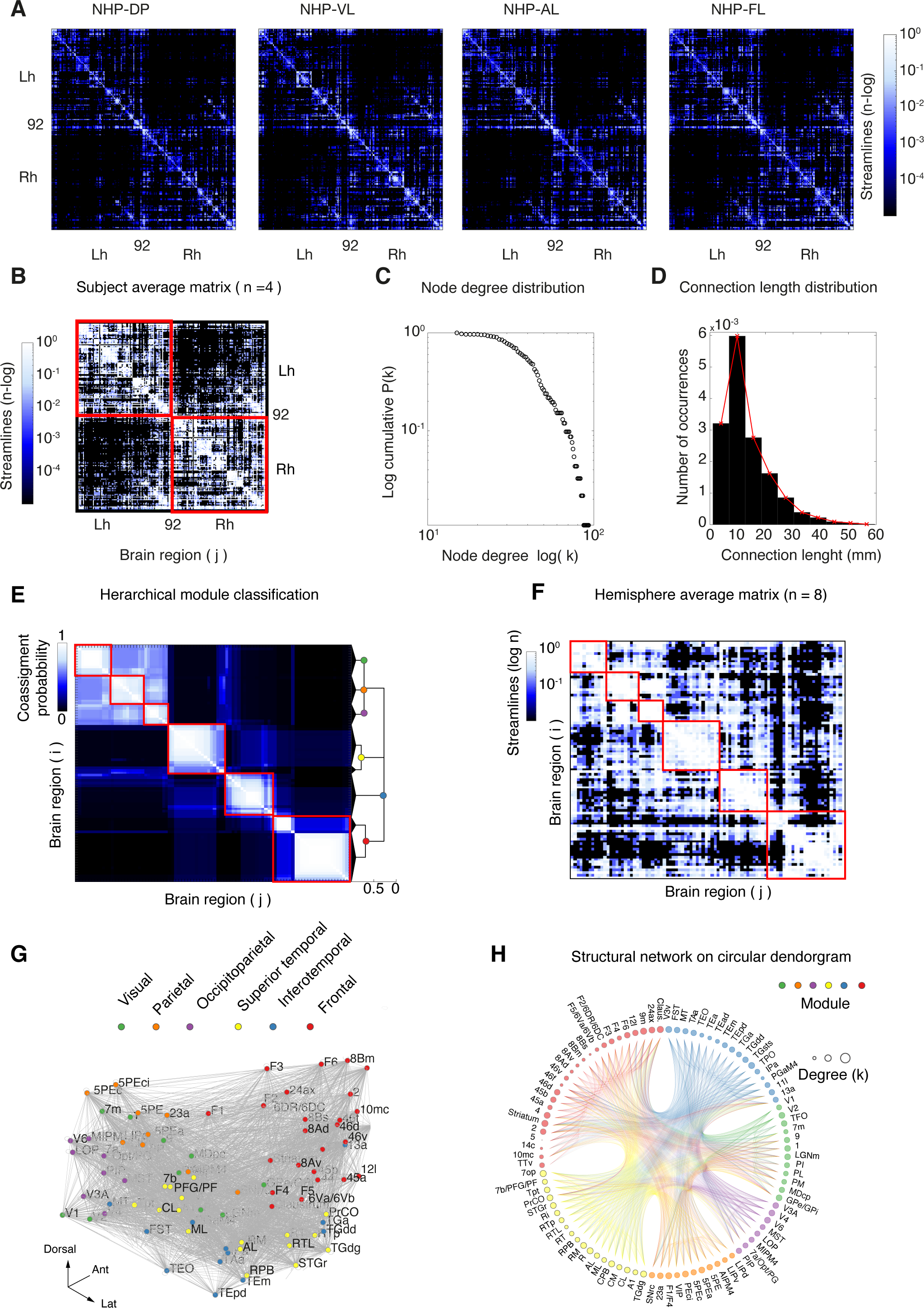
Organizing properties of structural networks. **A**. Structural connectivity matrices for each macaque monkey. **B**. Average structural connectivity showing the logarithmic number of streamlines touching every pair of ROIs for both hemispheres. **C**. Complementary cumulative distribution function (cCDF) of node degree shows a decrease in the probability of finding a highly connected done (*k*). **D** Distribution of path lengths shows a decay as a function of Euclidean distance. **E**. Co-assignment matrix and hierarchical dendrogram highlights module classification based on Q-value maximization. **F**. Average structural connectivity showing the logarithmic number of streamlines touching every pair of ROIs for both hemispheres. **G**. Modules plotted in anatomical space with nodes (visual, occipitotemporal, temporal, occipitoparietal, parietal, and frontal modules) color-coded according to their modular assignment. **H**. Circular dendrogram plot with hierarchical edge bundles aiding the visualization of the structural organization. The node face edge shows node degree connectivity.

From the low frequency (0.125 – 0.03) fMRI signals we computed the temporal correlation between the activity of each pair of brain regions over the whole time-series (e.g., static FVSNs). These correlations were constructed on weighted functional matrices (**Fig. 1C**). Given the existence of small non-zero values in the functional matrices, which may reflect the measurement of noise rather than the presence of an actual correlation (van Wijk et al., 2010), we applied a false discovery rate threshold (FDR p-value < 0.01) to the undirected adjacency matrices (Α_*ij*_) in order to determine which connections should be kept in the matrices,

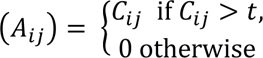

Using this threshold, the edges of (*C*_*ij*_) are a subset of the edges in (*A*_*ij*_). In subsequent steps we mostly used the Brain Connectivity Toolbox (Rubinov and Sporns, 2010) to calculate network measures on the corresponding matrices.

### 2.13 Construction of structural networks

To create the structural matrix the program 3dTrackID (Step 7) was used with DTI tensors, along with each pair of ROIs obtained from the D99 atlas. The output of 3dTrackID provides the connectivity matrix with the number of streamlines touching each pair of ROIs. For constructing Structural networks, we detect the number of streamlines (n > 10) that touch any pair of regions *i* and *j*. The logarithmically scaled number of streamlines were then arranged on a 184 x 184 connectivity matrix (*A*_*ij*_) of brain regions (**Fig. 1B**). We then binarized the networks for connectivity analyses.

## 3 Results

In the present study, we conducted two experimental free-viewing paradigm studies. Our first aim was to demonstrate the flexibility of the free-viewing paradigm for implementing both a classical analysis such as the general linear modeling (GLM) approach and those based on model free approaches such as coherence and independent component analyses (ICA).

### 3.1.1 Assessment of data using standard analytical techniques (Movie one)

Our first goal was to evaluate the overall brain activation pattern and to assess the effectiveness of the free-viewing approach. We took advantage of our improved implant design for functional imaging (Ortiz-Rios et al., 2018b) and mapped the BOLD response during the presentation of 30-seconds long movie clips. For GLM analyses we evaluate the presentation of movie versus baseline which we evaluated at a significant corrected threshold for false discovery rate (FDR q value) of < 0.05. Using this initial approach at the time of acquisition we quickly evaluated the quality of our data and confirmed significant activation (T-value colormap range 2.3 < 10, FDR q < 0.05) across the visual and higher-visual-related regions in all four macaque monkeys (**Supp. Fig.1A**). We then exploited the rich dynamics in the time series by implementing ICA analyses. ICA reveal the main free-viewing pattern related to the free-viewing BOLD modulation (30 secs ON and OFF periods) as the first main component explaining approximately 10% of the variance across run repetitions in each monkey (see **Table 2** and **Supp. Fig.2** for details). These preliminary analyses indicated that free-viewing elicited a strong driven pattern that could be revealed either by model oriented or model free analyses.

For the coherence analyses, the strength of the BOLD response was assessed on a voxel-by-voxel basis by first calculating the frequency-resolved power-spectrum and then determining the coherence between the BOLD peak frequency response (cycles/volume) to the stimulus repetition rate (0.016 Hz = 1/60 second). The movie scenes (n = 5) elicited a robust BOLD response modulation (mean ± SEM) across voxels in visual and high-order cortical regions in all four monkeys (see example modulation in the time course of cortical area MT of monkey DP, **Fig. 2B** and **Supp. Fig.5**).

**Figure 5.**
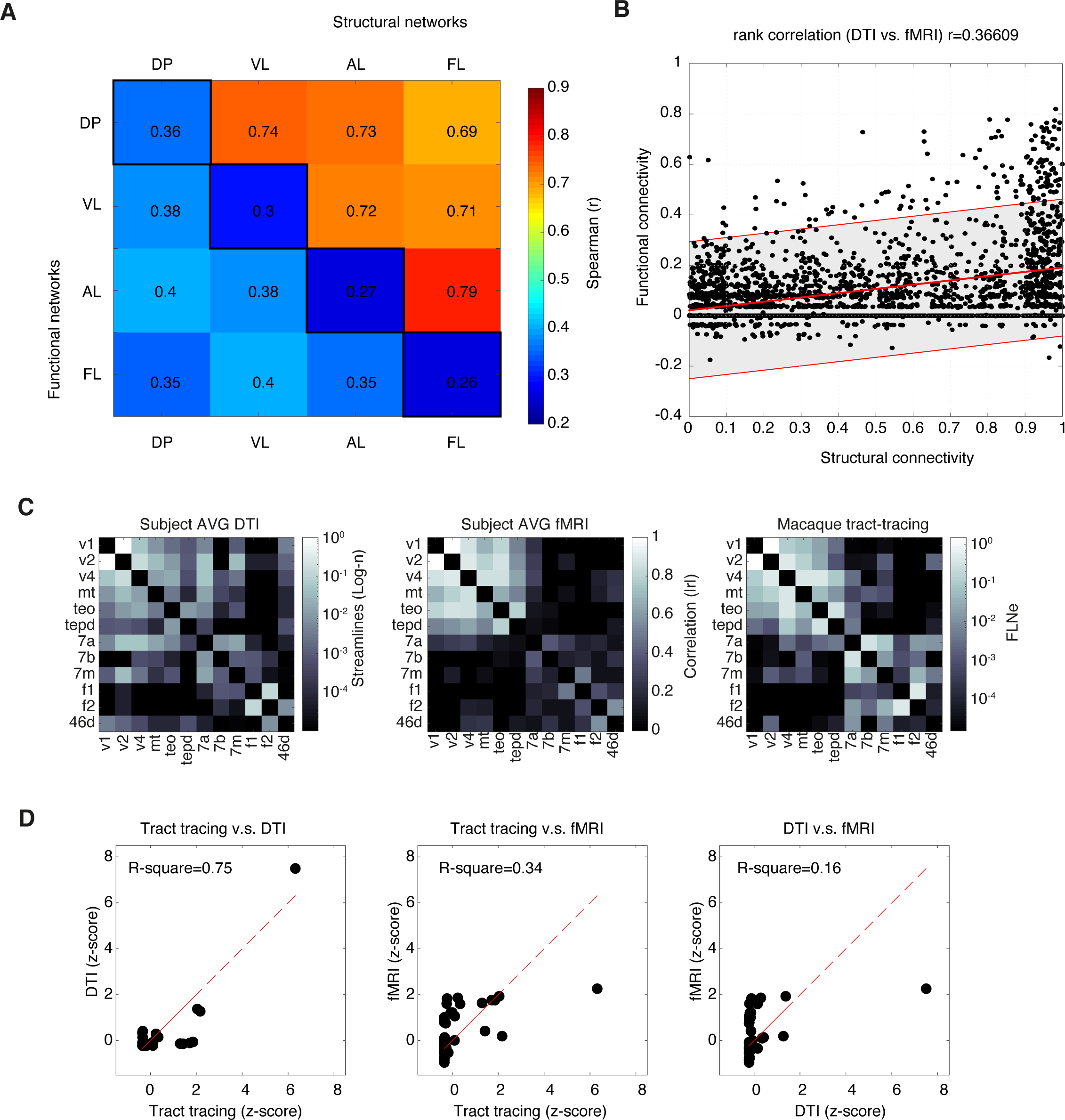
Comparing functional and structural networks. **A**. Confusion matrix between structural and functional networks showing Spearman correlation coefficients (r) between pairs of matrices. Along the diagonal (black squares) highlight the correlation within subjects across networks (e.g., free-viewing networks versus Structural networks of NHP DP). The lower left part of the matrix (light blue) shows the correlations across subjects for the free-viewing networks. The right top part of the matrix (orange red) shows the correlations across subjects for the structural networks. **B**. Scatter plot of correlation coefficients between structural networks and free-viewing networks with linear regression fit line (red) and 95% prediction interval (shaded grey area) showing a relatively low positive correlation (Spearman’s r = 0.37). **C**. Partial network matrices: DTI matrix (left), fMRI matrix (center) and macaque tract-tracing matrix (right). **D**. Scatter plots of z-scores values for the tract-tracing and DTI matrices are shown in the left panel. Spearman and R-square values are shown along with the linear fit (red dotted line). Similar scatter plots for comparisons of tract-tracing versus fMRI are also shown (center panel) and DTI versus fMRI (right panel).

In the visual and temporal cortex, we observed activation in V1, higher visual cortical areas (V2, V3, V4, V6), motion-sensitive regions (MT/MST/FST), and object-sensitive regions along the ventral stream (TEO, TE, TPO, TEm, IPa). Additionally, we also found bilateral activation in higher-order areas such as the frontal eye fields (FEF) and the lateral intraparietal area (LIP). These higher-level regions are known to be involved in the control of eye movements and in coordinating visuospatial attention (Buschman and Kastner, 2015; Gilbert and Li, 2013). This global cortical activation pattern was present in all scans and was consistent across all four macaque monkeys. Overall, our initial analyses (either GLM, ICA and coherence) confirmed the effectiveness of the free-viewing approach for eliciting reliable visual activation across the whole brain in all four macaques (**Fig. 2C**), as similarly observed in human brains cortical regions during movie watching (Hasson et al., 2004a).

#### 3.1.1.1 Centrality (Hubness) of free-viewing networks

While cortical mapping analyses largely revealed similar findings showing the regional activation of visual, higher visual, prefrontal and parietal cortices, the degree and involvement of cortical and subcortical and their interactions becomes more suitable to be evaluated and quantified using network graph analyses. Network analyses allow the detection of regional hubs that become more relevant than others during the viewing of visual scenes. Thus to overcome this challenge, we then constructed functional brain networks (**Fig. 1**) via individualized brain parcellations of D99 macaque brain atlas (Reveley et al., 2017). Thus, we next aimed to identify brain regions with high importance in Free-viewing networks by taking advantage of the recent developments on graph network analyses (Bassett and Sporns, 2017).

In order to find the most important – or central – regions in the network, we constructed static functional networks for each macaque monkey (n = 4; **Fig. 2D**). These networks were derived from low frequency (0.125 – 0.03) fMRI signals (see **Construction of free-viewing networks** in **Methods**) from which we then computed the temporal correlation between the activity patterns of each pair of brain regions (*i* and *j*). From each individual subject matrix we then computed the average connectivity matrix (**Fig. 2E**) and calculated eigenvector centrality (EC) for each brain region (Lohmann et al., 2010).

Eigenvector centrality (EC) measures the degree of a node influence in the network by scoring the node’s eigenvalue. A high eigenvalue of nodes indicates that the node is well-connected to many other well-connected nodes. For the thresholded matrix (*C*_*ij*_) let *x*_*i*_ be the eigenvector centrality of node *i* and *λ* largest eigenvalue and *x* the corresponding eigenvector. Eigenvector centrality is defined as,

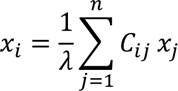

The proportional factor ^1^, is such that *x* is proportional to the sum of similarity scores of all connected nodes. We used the Brain Connectivity Toolbox to calculate eigenvector centrality. The MATLAB function is available at the Brain Connectivity Toolbox website (https://sites.google.com/site/bctnet/measures/list). In brief, EC measures the degree of a node influence in the network by scoring the node’s eigenvalue. A high-eigenvalue indicates that a node is well-connected to many other well-connected nodes. For the static networks we chose to highlight highly central regions with an eigenvector centrality value (EC) > 1x10^-4^ with a Z-score > 1.

Using this method, we identified a large-scale network engaged during movie watching (Z-score > 1 over the whole network; EC = 1x10^-4^) with the following functional hubs: MT/MST, V4, TEpd, IPa, Tpt, LIP, 8B, 46v, ProC, 12l, 13a, and striatum (**Fig. 2F**). Many of these regions are known to carry out specific visual functions related to object recognition, visual motion processing, eye movement control, and visual attention (Buschman and Kastner, 2015; Gilbert and Li, 2013; Russ and Leopold, 2015; Russ et al., 2016).

While many of the cortical areas (e.g., LIP, FEF, MT/MST, V4, TEpd, IPa) were also observed by contrast-based analyses; centrality measures of the same dataset enabled the identification of additional hubs that included regions (Tpt, 46v, ProC, 12l, 13a, striatum). These additional regions are less well delineated in terms of their contribution to visual function but could be identified from the connectivity analysis of free-viewing data. Importantly, half of the identified hubs (TEpd, LIP, 8B, 46v, 12l and 13a) were also previously identified from graph theoretical analyses of the macaque anatomical connectome as part of a ‘rich club’ of areas considered to be essential in facilitating global network communication (Harriger et al., 2012).

Overall, functional connectivity analyses revealed a widespread network of cortical and subcortical hub regions that were consistently present during free-viewing and that could only be partially identified with more conventional analyses. Next, we explored further the properties of Free-viewing networks.

### 3.1.2 Density and node degree of free-viewing networks

Previous network studies in both humans and monkeys have revealed that multiple network features are formed by the anatomical location and distance of regions within the brain (Hilgetag and Kaiser, 2004). Increase in local connectivity results in networks that prefer strong short-range connections which are thought to reflect intrinsic structural properties of brain connectivity that aim at reducing the metabolic cost of synchronizing activity across brain areas over the large-scale network (Watts and Strogatz, 1998).

Here, we first examine the connectivity matrices that evidently showed similar intra and interhemispheric correlation coefficients (**Fig. 3A**). In order to quantify these hemispheric properties, we calculated the network density (*ρ*) which is proportional and varies between zero and one, where *ρ* = 0 indicates no connection available, while *ρ* = 1 indicates that all possible connections exist, and 0 < *ρ* < 1 represents the fraction of all possible connections that are present in the network. The density from half of the matrix corresponding to the left hemisphere was similarly dense as the right hemispheres (Lh, μ = 0.16, ± σ = 0.01; Rh, μ = 0.21, ± σ, = 0.05). The density as measured over the whole brain (μ = 0.17; σ = 0.03) did not differ largely from the connectivity density of a single hemisphere alone, indicating the existence of similar functional properties across both left and right hemispheres. We further confirm this property by computing hemispheric correlations (Spearman’s r = 0.77, **Fig. 3D**). We quantify and summarize density measures for each hemisphere and macaque monkey in **Supp. Table 3**. These results indicated the existence of long-range connections present during free-viewing.

Next, we look at measures of node degree which aim at characterizing how correlations vary across regions (e.g., nodes). Node degree (*κ*) measures the number of edges (e.g., from correlation coefficients) of each node (*i*) with all other nodes (*j*) in the network. For our dataset average node degree (*κ*) ranged between 58 – 86 across subjects indicating the existence of nodes with a large number of connections (e.g., ∼ 70 > connections). To better characterized these results, we evaluated the cumulative distribution function (CDF) of node degree which showed a decay in the probability of finding a node (*κ*) with high degree (**Fig. 3B**, for each monkey and hemisphere see **Supp. Fig. 6**). The distribution decay indicated the existence of a large number of nodes with a smaller length connection and a smaller number of nodes with a larger length connection. From a network neuroscience perspective, these small number of nodes with many connections are considered essential in brain networks as their large-scale connectivity enables global and efficient communication within the network (Bassett and Bullmore, 2017).

**Figure 6.**
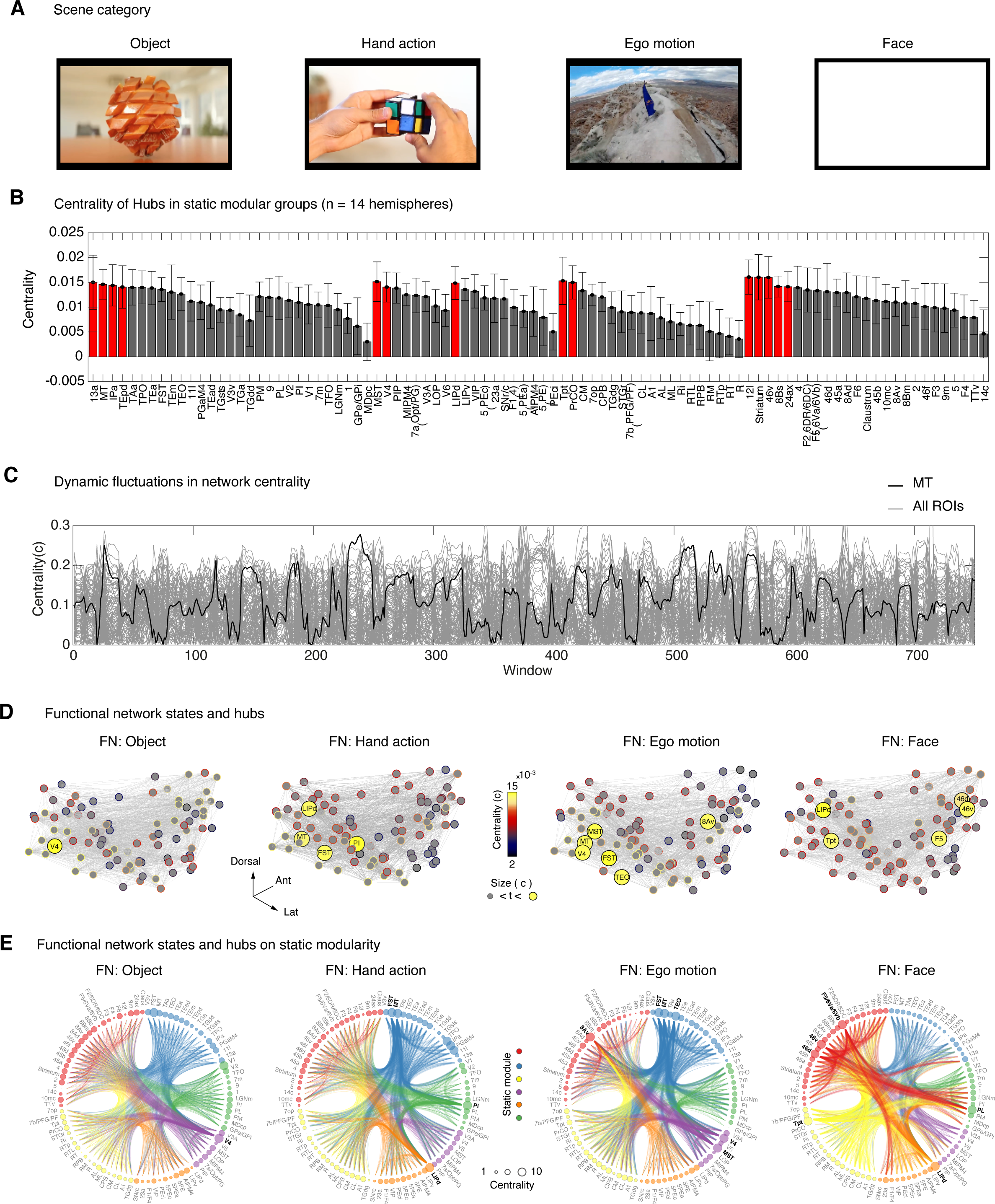
Dynamic reconfiguration of free-viewing network hubs. **A**. Four example image frames from the scenes presented to the NHPs during natural free-viewing. The presentation periods (30 seconds each) include object, hand action, ego-motion, and face scenes. **B**. Rank centrality averaged (mean +/- std) over 14 hemispheres show the consistency in hubs during free-viewing. Groups are shown for the static modular architecture obtained from structural networks. **C**. Dynamic fluctuations in network centrality. Gray plots show all ROIs centrality as a function of the sliding window. Black trace shows an example region (MT) with dynamic fluctuations in centrality during free-viewing. **D**. Eigenvector centrality (c) identify network hubs (c > 1x10^-4^; n for size <= 6) highlighting the central nodes during the presentation of each main categorical scene. The hubs change across the different scenes highlighting differential network states during natural free-viewing. **E**. Hierarchical edge bundling allows visualization of connectivity patterns in Free-viewing networks for each scene category on the underlying structural network modularity. The node size illustrates the overall network centrality value for all nodes with the threshold indicated on the black color of the node label. The edges that touch a highly central region show an increase in the edge thickness (size = 1).

### 3.1.3 Path length and small world topology of free-viewing networks

Next, we look at the characteristic path length (*L*), which measures the average shortest path for all possible pairs of nodes in the network. In brain networks, the Euclidean distance between local or clustered nodes tend to be short, thus regions that are anatomically nearby are therefore consider to be “economical” as their short distance minimizes metabolic cost, whereas the edges of long-range projections are fewer and sparse and are thought to have an increase integrative capacity over the network (Watts and Strogatz, 1998). The average path length for Free-viewing networks was short (μ = 2.21 mm, ± σ = 0.04, range: 2 – 2.5 mm,) and show greater differential distance as compare to their respective null-model rewired networks (*L* rewired μ = 1.8 mm, ± σ = 0.002, Free-viewing networks: 1.8 – 1.9 mm). **Supp. Table 3** and **Supp. Fig 9D** summarizes the characteristic path length for each hemisphere and macaque.

To further characterized path length measures, we plotted the distribution of path lengths and showed that the connection weight decays as a function of the Euclidean distance (node center of mass in mm). The distribution showed a peak distance around 15-20 mm (**Fig. 3C**, see **Supp.Fig.7** for each monkey and hemisphere) indicating the existence of strong long-range correlations. To further explore this feature of Free-viewing networks we plotted the connection weights (e.g., correlations coefficients) as a function of the Euclidean distance and found that on average the correlation decreases as a function of distance (**Fig. 3E**). To demonstrate the existence of high correlations among long-distance regions we separately plotted intra and interhemispheric correlation coefficients. Interestingly, short distance regions were more prone to be positively correlated but at longer distances positive correlations monotonically decreased. The majority of the weights across hemispheres were found beyond 20 mm with a peak at 40 mm, indicating the existence of long-distance correlations with anatomically distal related regions.

Furthermore, the characteristic path length (*L*) and the clustering coefficient (*Λ*) were used to determine the existence of small-world topology (σ) within the networks as compared to null-model rewired networks (Humphries and others 2006). Small-World network topology as indicated by the small world scalar show values greater than one (σ > 1 for Free-viewing networks, 3.3), path length greater than one ( *L* > 1 for Free-viewing networks, 1.2) and a clustering coefficient closer to one ∼ 1 (Free-viewing networks, 2.8). Overall, networks showed small-world network scalar consistent with previous graph theoretical analyses of human and animal brain networks (Bassett and Bullmore, 2017).

Overall, these findings suggested that the structural constraints from anatomical distance does not necessary capture the large-scale patterns of Free-viewing networks and that synchrony over long range distances may provide the network with an enabled functional architecture (Stiso and Bassett, 2018).

### 3.2 Bundle tractography of visual projections pathways

Functional networks are known to be highly dynamic and variable even during rest. Thus, in order to detect changes, it is ideal to first map the individual subject network from its static underlying structure. From structure we can then detect differences in network changes during free-viewing from which the dynamic changes in connectivity patterns could be inferred.

Towards this end, our next aim was to first evaluate whether the DTI data collected from awake monkeys could reveal the presence of prominent visual pathways. The macaque D99 atlas (Reveley et al., 2017) was used to map each macaque brain individually using its own native space (e.g., within a scan session). From diffusion-weighted analyses, we computed eigenvalues, eigenvectors, mean diffusivity (MD) to obtain fractional anisotropy (FA) map (**Supp. Fig. 1B**) (Basser and Pierpaoli, 1996). We then identified white matter tracts - streamlines - of the occipital cortex using a pair of regions selected from the warped D99 macaque-atlas. Streamlines projections passing between two pairs of ROIs were considered members of the same white matter bundle (see **Methods** for details). We then computed diffusion tensors using multi-directional tractography estimated the tensor uncertainty (Taylor and Saad, 2013b; Taylor et al., 2012).

The streamlines from the right LGN to the right V1 in all four macaques (**Supp. Fig.5**) showed similar bundle tracks highlighting the geniculostriate pathway; also known as the optic radiation (OR). The OR bundle contains neuronal projections that carry visual information from the retina to visual cortex. Since our stimulation consists of visual scenery, the OR evaluation becomes relevant for our functional connectivity analysis. The OR bundle as measured by DTI projects mostly posteriorly towards V1 of the same hemisphere and shows streamlines mostly colored in green (e.g., anterior-posterior direction). In **Supp. Fig. 5**, we show additional visual bundle tracts (e.g., the forceps major) crossing over hemispheres of each monkey.

### 3.3 Density and node degree of structural networks

Having confirmed the sensitivity of our DTI setup to reveal sensible anatomical projections, we then constructed Structural networks by detecting the number of streamlines (n > 10) that touch any pair of regions (*i*) and (*j*). The logarithmically scaled number of streamlines were then arranged in a 184 x 184 weighted undirected (symmetrical) connectivity matrix (Α_*ij*_) of brain regions (**Fig. 4A**) with the same size and regional organization as implemented for Free-viewing networks. Given the structural similarity across the matrices for each monkey (**Fig. 4A**) we then average the structural matrices and obtain an average structural matrix (**Fig. 4B**). We then calculated the density of Structural networks and found higher connection density (*ρ*) within hemispheres (Lh, μ = 0.42, ± σ = 0.03; Rh, μ = 0.40, ± σ = 0.04) as compared to the total density when sampling both hemispheres (both; μ = 0.28, ± σ = 0.02). Such smaller connection density across hemispheres as compared to within hemispheres largely reflects the smaller number of large distance streamlines passing through the corpus callosum as similarly found in human DTI-based structural networks (Bonilha et al., 2015).

We then evaluated the number of edge connections to each node or node degree (*ρ*) distribution which shows a decay in the probability of finding a node with high node degree (see **Fig. 4C** for average network and **Supp. Fig. 8**) for each independent hemisphere and network class; (*κ*) range across macaques, Structural networks (*κ*): 148 – 159).

From anatomical tract-tracing (Markov et al., 2014) and graph theoretical studies (Song et al., 2014) is it well-establish that brain regions are highly and densely connected with proximal neighboring regions. By measuring the clustering coefficient (*C*), we could quantify the number of connections that were present between the nearest neighbors of a node as compare to the total number of all possible connections. Clustering coefficients were compared with those obtained from random networks which are known to have lower average clustering as oppose to brain networks which show higher clustering (Bullmore and Sporns, 2009). For our binarized Structural networks the clustering coefficient *C* shows relatively higher values (see **Supp. Fig. 9A**, NHP range: 0.69 – 0.71; μ = 0.7, ± σ = 0.01) as compare to rewired networks with the same degree distribution under the null-hypothesis (NHP range: 0.45 – 0.52, μ = 0.5, ± σ = 0.69 x 10^-3^). Overall, both node degree and clustering coefficient analyses indicated that the resultant properties we obtain from DTI-based structural networks could enable the global and efficient communication within the network (Hilgetag and Kaiser, 2004; Kaiser and Hilgetag, 2006).

#### 3.3.1 Path length and small-world topology of structural networks

Next, we look at characteristic path length (*L*), which measures the average shortest path for all possible pair of nodes in the network. The characteristic path length (*L*) of Structural networks measured from the binary graphs shows a classical shorter path length (*L* range across NHPs, Structural networks: 1.73 – 1.8 mm, μ = 1.76 mm, ± σ = 0.001) as compare to the null-model rewired networks (see **Supp. Fig. 9B**, *L* rewired Structural networks: 1.7 1.75 mm, μ = 1.72 mm, ± σ = 0.4 x 10^-5^). We then calculated the path length distribution (**Fig. 4C)** of the average SN which shows a peak between 10 -12 mm consistent with the peak distance of anatomical tract-tracing studies of the macaque (Markov et al., 2014). This path length distribution was also consistent with interareal distances across the cerebral cortex of multiple species (Song et al., 2014).

Furthermore, the characteristic path length (*L*) and the clustering coefficient (*Λ*) were additionally used to determine the existence of small-world topology (σ) within the networks as compared to null-model rewired networks (Humphries and others 2006). Small-World network topology as indicated by the small world scalar show values greater than one (σ > 1 for Structural networks; 1.98; Free-viewing networks, 3.3), path length greater than one ( *L* > 1 for Structural networks, 2; for Free-viewing networks, 1.2) and a clustering coefficient closer to one ∼ 1 (for Structural networks, 1.02; Free-viewing networks, 2.8). Overall, networks showed small-world network scalar consistent with previous graph theoretical analyses of human and animal brain networks (Bassett and Bullmore, 2017).

In summary, Structural networks show differential connectivity structure as compared to Free-viewing networks. The topological properties of the networks showed similar organizing principles as those described in other species and from derivation using other techniques (Song et al., 2014). From the partial connectivity comparisons these analytical findings also indicate that structural networks as derived from DTI could capture the overall structural topology (Hilgetag and Kaiser, 2004; Kaiser and Hilgetag, 2006; Markov et al., 2014).

### 3.4 Modularity of static-structural networks

From the average structural matrix we then aimed at identifying network modules; groups of regions that are densely connected among each other but sparsely connected across groups (Sporns and Betzel, 2016). To identify network modules we used modularity maximization; a data-driven approach for hierarchical clustering of nodes into network modules (Rosenthal et al., 2018). We calculated network modularity (*Q*) via a hierarchical consensus algorithm. (*Q*) maximization is accomplished by comparing the adjacency matrix (*A*_*ij*_) with the expected null connectivity model (*P*_*ij*_), where larger (*Q*) values are indicative of a higher quality. The data driven approach allows for the grouping of nodes into modules that show high internal density as would maximally be expected from the null model. Consensus modularity (*Q*) is defined as:

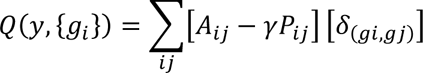

Where *γ* is the resolution parameter used for optimization; *g*_*i*_ϵ{1, …, *C*} is the module assignment of node *i*, where the Kronecker delta function δ_(*gi*,*gj*)_ equals one, if nodes *i* and *j* belong to the same module (*g*_*i*_ = *g*_*j*_). The function ensures the total weight of within-module edges is less than that of the null model. We used the multiresolution consensus function to calculate modularity (Jeub et al., 2018) (https://github.com/LJeub/HierarchicalConsensus).

Using this method, we found six hierarchical modules partitions that overall captured the well-known macaque brain organization (**Fig.4E** and **F**). The organized brain structure is evident on the 3-dimensional anatomical network plot (**Fig.4G**) where regions preserved their center of mass and anatomical coordinate. We identified visual, occipitotemporal, temporal, occipitoparietal, parietal, and frontal modules. Some of the modules show the ability to further subdivide within the hierarchy (e.g., frontal module). For our Structural networks, we show subdivisions up to level six which overall recapitulated the structural and functional brain connectivity of the macaque monkey (Harriger et al., 2012). To ease the visualization of the connectivity structure among modules, we used hierarchical bundling of the edges (Holten, 2006) and reduced visual clutter. The bundle edge technique allows mapping the implicit adjacency across modules and connectivity within regions. Additionally, we mapped, the number of connecting edges to each node or node degree (*κ*) on the node face size, allowing us to represent the overall brain connectivity within a single graph.

Overall these results recapitulate the well-known structural and functional connectivity of macaque brain networks (Harriger et al., 2012) and are in accordance to previous macroscale mapping of human anatomical brain networks derived from DTI structural data (Power et al., 2011).

#### 3.4.1 Network comparisons: Free-viewing networks and Structural networks and macaque tract-tracing networks

To assess the similarity between each macaque network or intra-subject correlation, we calculated the Spearman correlation coefficient (*r*) between any pairs of matrices and found consistent relationships between individual pairs of macaque structural networks (see upper diagonal matrix in **Fig. 5A**, Structural networks range (*r*) = 0.60 – 0.79). To further explore the relationships among Structural networks, Free-viewing networks we first evaluated the correlations between whole brain undirected Free-viewing networks and Structural networks. These comparisons show relatively low correlations (range, (*r*) = 0.2–0.36) across networks (e.g., Free-viewing networks versus Structural networks) within each macaque subject (see lower diagonal of confusion matrix on **Fig.5A**). The correlation coefficients between Structural networks and Free-viewing networks showed relatively low positive correlation (Spearman’s r = 0.37, linear fit 95% prediction interval). To compare networks with tract-tracing studies we match the ROI label nomenclature of the regions in the Markov study (F99 atlas) to our present study (D99 atlas). We found twelve matching ROI regions that were then used to compare networks across modalities. The average partial matrices (**Fig. 5C**, see **Supp.Fig.11** for each monkey partial matrices) were z-scored within-subjects and then averaged for comparisons to the z-score tract-tracing matrix. The R-square correlation between tract-tracing and DTI showed relative higher value (R^2^ = 0.75) as compared to tract-tracing versus FV-fMRI (R^2^ = 0.34) or DTI versus FV-fMRI (R^2^= 0.16) indicating a higher correspondence between macaque anatomical networks (**Fig.5D**).

### 3.5 GLM contrast analyses of movie sequence two

A previous naturalistic viewing study in macaques extracted basic visual features, such as luminance, motion, and faces from visual scenes (Russ and Leopold, 2015), and then tested the relationship between the occurrence of each feature with the ongoing BOLD response. Here we extended this approach by creating purpose-build movie scenes with the following categories: object, hand action, ego-motion, and faces (**Supp.Fig. 12A**). We then contrasted each movie scene with their respective frame-by-frame matched controls (optic-flow, phase scramble, salience-contour, and tile scramble). Using contrast analyses, we found distinct – but largely overlapping – patterns of activation along the superior temporal sulcus (STS) for each movie scene (see **Supp.Fig. 12B** for example monkey DP, q FDR < 0.05, p < 1.7^−5^, cluster size > 10 voxels; and **Supp.Fig. 12C** for example monkey AL, q FDR < 0.05, p < 8.5^−6^, cluster size > 10 voxels). Across all scenes, we found consistent activation in visual areas V2, motion-sensitive area MT and ventral stream areas V4 and TEO. A closer inspection of the original scene conditions revealed specific activation patterns for each scene, which we describe in greater detail in the supplementary material and table section for details.

#### 3.5.1 Time-resolved connectivity of free-viewing networks

The previous analyses on Structural networks allow us to identify the static modular structure of macaque brain networks. This static structure could be used to detect networks changes during natural free-viewing. With this aim in mind we first quantify some of the structural network properties and compare structural, functional, and partial anatomical networks derived from macaque tract-tracing studies (Markov et al., 2014).

Time-varying functional connectivity was estimated using the multiplication of temporal derivatives (MTD), which calculates each sample product of temporal derivatives for pairwise time series (Shine et al., 2015). For each time point *t*, the MTD is defined for the pairwise interaction between region *i* and *j*,

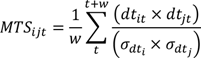

Where *dt* is the first temporal derivative of either *i* and *j* at time *t*, and *σ* is the standard deviation of temporal derivative for time series *i* and *j*, and *w* is the window length (n samples = 12 x 1.5 TR = 18 sec) of the moving average. See coupling for more details (https://github.com/macshine/coupling/).

Since several regions showed specific scene-content dependent activation, our next aim was to understand how functional interactions between areas might reorganize across the different movie epochs. To this end, we built on our previous analyses and calculated network centrality for all regions during a moving time window (**Fig. 4A**).

The overall network showed dynamic changes in network centrality throughout the entire scanning period (see example fluctuations in centrality from area MT). Interestingly, while the overall global centrality pattern largely reflects the induced sensory modulation, we observed that many regions significantly vary in their centrality across both movie period and darkness period. From the specific scene epoch matrix, we calculated eigenvector centrality (Z-score > 1, c > 1 x 10^-4^ for n for size <= 6) and identified network hubs for each main scene category (**Fig. 4B**).

As a first instance, we observed changes in network centrality across all four original movie scenes. For a more detailed visualization of the network interactions for each scene, we then plotted the resulting Free-viewing networks in a circular dendrogram with the node face size representing degree centrality. From the modularity of structural networks of each animal (see **Modularity of static structural networks**) we derived the color code classification for each brain region in the group. The edge-width highlights the major projection bundles between hubs in the network, while the widths of all other edges were kept constant (**Fig. 4C**). In what follows, we describe the main functional connections for each network state and relate these to existing knowledge about macaque anatomical connectivity of the hub regions identified using network centrality.

For the object network, we observed high network centrality for area V4 (in the occipitotemporal module color-coded in purple). The majority of connections to or from the V4 bundle diffuse into prefrontal (red) and superior temporal (yellow) modules. Area V4 is well-known to be involved in visual object recognition (Roe et al., 2012), which is confirmed by the high network centrality we observed during the object scene condition. The connections bundles we observed in our Free-viewing networks between V4 and FEF were also consistent with the reciprocal connections between both areas (Markov et al., 2014) and the proposed importance of this connection in visuo-saccadic planning and attention (Roe et al., 2012).

For the hand-action network, we found high centrality in regions MT, FST, LIPd, and subcortically in the inferior pulvinar (PI) nucleus. Regions MT, FST show projection bundle-tracts that connect to all five network modules with particular convergence in areas LIP in the parietal module (orange) and to the PI nucleus (visual module in green). Area LIP is involved in visuospatial attention (Buschman and Kastner, 2015) and the spatial coordination of limb movements (Gottlieb, 2007; Oristaglio et al., 2006), which is in line with our results for the hand-action network. Further, we identified functional connections between the inferior pulvinar and area MT/MST; an anatomical connection previously described from tract-tracing studies in the macaque (Boussaoud et al., 1990; Kaas and Lyon, 2007). Moreover, dorsal motion areas are known to be connected with the posterior parietal cortex, including area LIP (Boussaoud et al., 1990; Markov et al., 2014), supporting our connectivity findings related to our hand-action network.

For the ego-motion network, we observed hubs MT, FST, TEO and V4 showing functional connections with frontal lobe region such areas 8av (e.g., FEF region). This network showed stark differences in comparison to the object network. The object network showed only area V4 as the main central region. Anatomically, it is well established that areas V4, TEO, MT, FST, MST are inter-connected linking information processing between dorsal and ventral visual pathways for motion analyses (Boussaoud et al., 1990). Further, the centrality of motion complex regions in the STS confirms that motion sensitive regions became essential during increasing levels of visual motion. Additionally, our findings with the ego-motion scenes recapitulate recent fMRI studies in macaques, indicating a prominent visual motion drive during free-viewing of dynamic visual scenes (Russ et al., 2016).

For the face network, we found hubs in areas LIP, Tpt, 46, ventral premotor cortex (e.g., areas F5/6Va/6Vb) and subcortically the lateral pulvinar nucleus (PL). These results indicated that our centrality measures were able to capture additional important features of the interareal interactions not available with standard GLM contrast analyses. On the network, we observed a drastic shift in connectivity in comparison to those obtained with other scenes. Specifically, face networks within the frontal module showed increases in connectivity locally and with large bundles connecting with regions in the superior temporal module (area Tpt), parietal (area LIP), and visual (lateral pulvinar) modules. The lateral pulvinar is known to project to early visual areas of the ventral cortical stream (Kaas and Lyon, 2007). Additionally, it receives projections specifically from cortical face patches in the temporal lobe (Grimaldi et al., 2016). Moreover, the ventrolateral and ventral premotor cortex is connected to regions in the superior temporal and parietal cortex (Petrides and Pandya, 2009), a pathway that is thought to integrate social facial gestures (Shepherd and Freiwald, 2018) with vocal sounds (Ortiz-Rios et al., 2015). Our centrality state during free-viewing of face scenes further implicates this network in social communication (Sliwa and Freiwald, 2017) as similarly found in humans (Lahnakoski et al., 2012).

Taken together, our connectivity analyses allowed us to observe changes in network interactions across cortico-cortical and thalamocortical network pathways, with some interactions which were not evident from traditional mapping analyses. The dynamic changes we observed during free-viewing highlight the importance of the ongoing motor and cognitive processes that take place during more naturalistic experimental conditions.

## 4 Discussion

In this study, we used a free-viewing paradigm in combination with graph theoretical analyses to investigate how functional networks reorganize during active natural vision. Our analyses with varying functional connectivity revealed specific thalamo-cortical and cortico-cortical functional interactions that reconfigured depending on the presence of objects, motion, face and actions within the scenes. Overall, across monkeys and movie scenes we found a consistent functional network engaged during free-viewing that included hub regions in frontal (FEF), parietal (LIP, Tpt), and occipitotemporal cortex (MT, V4 and TEpd) among others.

In the following section we discuss our results with respect to the current knowledge of the anatomical connectivity between hub regions and the involvement of these regions in visual cognitive processes. Additionally, we discuss how naturalistic imaging could be exploited further for the study of functional brain networks at large. Lastly, we envision how new paradigms, in combination with networks neuroscience, can help unravel new insights into the spatiotemporal dynamics of large-scale neuronal networks.

### Network state configurations during natural free-viewing

One fundamental principle of brain organization is the existence of functional hubs that enable efficient neuronal communication and the integration information over long distances in the network (van den Heuvel and Sporns, 2013). By applying graph theoretical methods, we were able to identify regions beyond the traditional visually engaged areas. These regions included multimodal areas (Tpt, 46v), prefrontal cortex (12l, 13a, ProC), pulvinar and the striatum. Anatomically, these hubs are known to contain widespread connections, but with so far largely unappreciated contributions to natural vision.

When we focused the analysis on movie scenes that contained faces, we found hubs in parietal (LIP), superior temporal (Tpt), prefrontal (area 46), premotor (F5), and subcortically in the lateral pulvinar nucleus. The dorsal network that connects premotor regions with the superior temporal and parietal regions (Petrides and Pandya, 2009) is known to integrate social facial gestures (Shepherd and Freiwald, 2018) and vocal sounds (Ortiz-Rios et al., 2015). The hubs we found in frontal and premotor cortex (e.g., 46 and F5) along with the superior temporal (Tpt) region suggests that the free-viewing of faces engaged this network for the extraction of higher-order social features within the scenes (Shepherd and Freiwald, 2018). The projections from the superior temporal region (Tpt) and parietal cortex (LIP) may provide multimodal and spatial information to face sensitive regions in the IT cortex (Cusick et al., 1995; Seltzer and Pandya, 1994). Subcortically, we also found high centrality of the lateral pulvinar (PL) during the face scene periods. The PL anatomically projects to the inferotemporal cortex (Kaas and Lyon, 2007) and receives projections specifically from face patches (Grimaldi et al., 2016). An additional interesting point relates to the medial pulvinar (ML) which projects to the anterior STG and functionally is involved in higher-level features of vocal sounds. Both thalamic nuclei (medial and lateral pulvinar) might be a source for the modulation of both vocal and facial information at the level of STS (Bruce et al., 1981; Grimaldi et al., 2016; Scott et al., 2017; Smiley and Falchier, 2009).

Our analysis also revealed the recruitment of a different network engaged in the processing of hand action scenes. For these, we observed specific connections with areas LIP, MT/MST, and the inferior pulvinar (PI). Regions in the dorsal stream (e.g., LIP, MT/MST) are well-known to coordinate visuospatial information from eye and limb movements (Gottlieb, 2007; Oristaglio et al., 2006). Moreover, these areas receive projections from the inferior pulvinar (Boussaoud et al., 1990; Kaas and Lyon, 2007), consistent with the observed connectivity patterns for the hand-action network. In the cortex, we found regions FEF, LIP, 46, 13, 24, F5, and TE as network hubs, in line with previous graph-theoretical analyses of macaque anatomical networks (Harriger et al., 2012). Area TE, corresponds to the well-known AF face patch region of the inferotemporal cortex (Tsao et al., 2003) and their neurons (Tsao et al., 2003) have been recently found to response to the overall spatial layout of natural visual scenes (McMahon et al., 2015). Additionally, area TE responsed to complex biological actions and received projections from the overlying polysensory STS region (Bruce et al., 1981) and parietal cortices which might provide TE regions with multimodal spatial information.

Additionally, we also found recurrent network hubs, such as the motion complex network (e.g., areas MT/MST/FST) that emerge as central during the presentation of multiple visual scenes. Not surprisingly, the emergence of the motion complex network during free-viewing indicates the necessity for processing a substantial increase in visual complexity and visual motion in our selection of visual scenes. These findings are also in accordance with recent macaque fMRI studies using naturalistic stimulation that investigated the influence of self-induced and visual motion (Russ et al., 2016) and found a strong dominance of visual motion in the fMRI activation patterns along the STS.

Similarly to MT, we also found high centrality in area V4, which is known to be involved in visual shape and object recognition (Roe et al., 2012) and for our object scene condition, it follows the role of area V4 in processing objects features. The object network showed stark differences in comparison to the ego-motion network. The ego-motion network shows an engagement beyond V4 and included motion complex regions (MT/MST FST). Additionally, area V4 is known to play a role in visuospatial attention (Moran and Desimone, 1985; Roe et al., 2012), and to receives inputs from and project to the FEF region (Markov et al., 2014) in accordance to our findings showing high centrality for area FEF. The connections we observed in free-viewing networks confirms the role that area FEF and area V4 play in top-down visuo-saccadic functions (Ekstrom et al., 2009; Moore and Armstrong, 2003; Roe et al., 2012). Additionally, object motion related regions (e.g., V4, MT, MST) became functional hubs along with saccade-related regions (e.g., LIP and FEF) indicating that, in general, movie viewing particularly engages this network configuration. Moreover, these results are in close accordance with previous ego-motion fMRI studies in macaques (Cottereau et al., 2017) showing the involvement of areas MST and FEF during control ego-flow motion conditions.

It is well established that areas LIP and FEF are involved in saccadic eye movements and the visuospatial guidance of eye movements and attention (Bisley and Goldberg, 2010; Buschman and Kastner, 2015). We refer to this overall network configuration as a visuo-saccadic network which requires higher-level visual object processes and motion analyses (visual, e.g., V4 and MT/MST) and is also concerned with extracting and guiding the visuospatial coordination of eye movements (e.g., LIP and FEF) during free-viewing. Single-unit studies of macaque FEF during natural visual search indicates that FEF activity might contribute to top-down selection (Glaser et al., 2019; Juan et al., 2004; Phillips and Segraves, 2010; Ramkumar et al., 2016), while neurons in area V4 might converge bottom-up salient visual information with top-down selection of eye-movements target locations (Mazer and Gallant, 2003; Zhou and Desimone, 2011). Moreover, similar visuo-saccadic networks were also reported in marmosets and humans during free-viewing (Schaeffer et al., 2019) indicating a conservation of this network across primate species.

Overall, our findings suggest that free-viewing networks captured a large part of the underlying anatomical and functional architecture (Kaiser and Hilgetag, 2006; McMahon et al., 2015). Moreover, the robustness of the visuo-saccadic network and the high feasibility of detecting it during free-viewing (e.g., no task based training inside the scanner) makes it an ideal network for studying dysfunctional changes in patients with Parkinson’s (Fukushima et al., 1994) and other neurodegenerative diseases (Antoniades and Kennard, 2015) often expressing deficits in saccadic function.

### Comparisons of free-viewing and structural networks

The relationships between structural and functional networks in NHPs is a critical step for developing large-scale neural simulations and informative models of brain function (Shen et al., 2019). Our initial work here on network analyses demonstrates the feasibility of obtaining meaningful structural graphs derived from awake imaging data. Importantly, our analyses also showed the existence of well-known network features such as a decay in node degree distribution, local clustering and short-path length; all characteristic properties of small-world network architecture (Watts and Strogatz, 1998).

Comparisons across networks using linear regression showed a relatively low positive correlation (Spearman’s r = 0.37), recapitulating previous comparisons across structural and functional networks in humans (Honey et al., 2009). The correlations we found within NHP networks (e.g., same subject FVN and SN) ranged between (r range; 0.26 – 0.36) indicating fundamental differences in network architecture; also evident from the connectivity matrices (**Figure 2D** and **Figure 4A**). Correlations within subject structural networks were relatively high (r range; 0.69 – 0.79) as compared within subject free-viewing networks (r range; 0.35 – 0.4). Interestingly, for free-viewing networks we observed and increased connection lengths as compared to structural networks (**Figure 3C** and **Figure 4D**). When examining the correlation coefficients of free-viewing networks as a function of connection distance we observed a small number of high correlated nodes with long path length distribution (**Figure 3E**). These long-distance connections might result from synchronous time-series among poly-synaptic distal regions suggesting that during free-viewing functional interactions might be less distance-dependent than structural connections (Bettinardi et al., 2017). Empirically, NHP structural connections are known to be constrained by the embedded brain architecture and to carry metabolic cost that generally reflects a decrease in the probability of finding a long-range connection

(Horvát et al., 2016). In our structural networks, such local clustering was also evident by the hierarchical modular structure which provided classifications largely reflecting lobular organization (**Figure 4G**). More importantly, our approach allows us to study the free-viewing networks departing from a static architecture as a way to infer edge divergence and changes in hub centrality.

### Advantages, limitations and future directions for free-viewing networks

In primates, most macroscale functional mapping studies have focused on the existence of intrinsic functional networks that show similar properties as those found in humans’ resting-state networks (Vincent et al., 2007). resting-state networks are known to be highly variable across individuals and evolving periods of an imaging session (Hutchison et al., 2013; Nikolaou et al., 2016; Preti et al., 2017). As a result, under resting-state conditions, it is challenging to predict the internal states that could trigger dynamical shifts in functional connectivity across any given time window and individual.

Naturalistic imaging in NHPs provides similar advantages as those previously demonstrated in human neuroimaging studies (see Eickhoff et al., 2020 for review) that are complementary to RSN: (1) the ability to relate networks structure with the external stimulation (e.g. movie) sequence paradigm, (2) the ability to consistently drive distal and higher-order regions within the network (e.g. cognitive states) (3), the ability to induce reliable activation patterns across individuals (e.g. high inter-subject correlations) (4) and an increase in contrast-to-noise ratio (e.g. improve data quality). Additionally, naturalistic imaging engages NHPs during movie watching, possibly comforting animals during scanning periods, and results in reduced movement artifacts. In our data, we experienced minimal motion deviations (e.g., shifts below 0.5 mm and rotations below 0.5 degrees) which resulted in an effective motion-free time-series (see example time course in **Fig.2**). Furthermore, the activation patterns we obtained across subjects and replicable centrality measures highlight the effectiveness and reliability of the naturalistic imaging approach to induce similar effects across subjects. The fact that we obtained our data within two repetitions of a five-minute scan, demonstrates to the high contrast to noise ratio available in our dataset, additionally evident from the temporal SNR analysis.

Most importantly, naturalistic imaging allowed us to capture network dynamics under more complex, ecologically valid states. In our study, we created free-viewing networks from which we captured distinct thalamocortical and cortico-cortical interactions that were not readily observable using more standard analyses. However, it is important to note that while we linked network interactions with the content of the movie, our study is limited by the epoch of presentation rather than by the evolving structural temporal motifs in networks (Sizemore and Bassett, 2018). Another limitation relates to the variation of movie content that we presented. It is conceivable that a wider range of scene content presentation might have further refined the implicated network pattern. For example, the study by Hasson et al., 2008 showed that higher-level regions (such as FEF, TPJ and STS) showed disruptions in temporal structure during scramble scenes while] motion sensitive regions ( MT+) were insensitive to temporal disruptions. An interesting future approach for measuring network changes in temporal structure manipulations will be to create gradual and slow changes in scene content (either in forward or backward manner) as a strategy to pick up and down shifts in states of the network.

Another avenue of interest for exploiting the dynamics of naturalistic networks lies in the concurrent recording of eye movements. The study by (Russ et al., 2016), specifically quantified the effects of saccadic eye movements on functional activation maps by means of regression analysis. Their study demonstrated that the variation of eye-movements resulted in very similar eye-movement patterns across repetitions of the same movie. From our datasets we expect that differential eye movement patterns might have contributed to the networks states we observed during more ecological valid conditions as compared to fixation-based tasks. Furthermore, on a more technical note, in our study we sampled functional data at a 1.5 second per scan, limiting our ability to infer network effects from multiple instances of saccades events occurring during a movie viewing epoch. Moreover, eye-movement recordings were not always successful on all the animals, limiting our ability to relate brain networks with intrinsic eye-movement data. The eye-tracking apparatus on the vertical-scanner had been removed on every scanning session limiting our ability to maintain pre-calibrated data. In the future, it will be of interest to further explore the link between network dynamics and eye movements, including pupil dilation, which is known to cause fluctuations in arousal states that involve cholinergic modulation of large-circuits (Shine et al., 2016). Together with the application of more advanced network analyses, it will further aid the detection of recurrent temporal network states in NHPs (Bassett et al., 2011) during free-viewing.

We believe that the approaches outlined here in NHPs will allow disentangling spatiotemporal dynamics that might emerge as repeatable network states during cognitive processing. Multimodal imaging in combination with large-scale neurophysiological recordings this approach will provide an important window into the underlying network mechanisms of cognition. With the expansion of graph-theoretical tools, a robust computational framework has emerged in neuroscience (Bassett and Sporns, 2017; Bullmore and Sporns, 2009) which has led to dramatic advances in our understanding of the principles guiding the organization of brain networks. In the future, the use of network analysis tools in combination with optogenetic or chemogenetic interrogation techniques (Ortiz-Rios et al., 2018a) might enable the targeted control of neural circuits for the manipulation of global network states.

### Code Accessibility

Imaging data reported in this paper have been shared on and are additionally available in https://github.com/ortizriosm/natural-vision-connectivity along with data pre-processing scripts. Additional information about non-human primate research can be found on the website of SchmidLab (https://research.ncl.ac.uk/schmidlab/). Additional information about brain connectivity research can be found on the website of the Dynamic Connectome Lab (https://www.dynamic-connectome.org).

## Acknowledgments

We wish to thank Ian Milne and Joe Wardle for their help in developing hardware and equipment for MRI-compatible imaging. We would also like to thank the CBC staff for their constant effort with animal training and handling during experiments. Marcus Kaiser was supported by Wellcome Trust (102037), Engineering and Physical Sciences Research Council (NS/A000026/1, EP/N031962/1), and the Guangci Professorship Program of Ruijin Hospital (Shanghai Jiao Tong Univ.) This work was funded by ERC OptoVision 637638 to Michael Schmid.

---

**Supp. Fig. 1.**
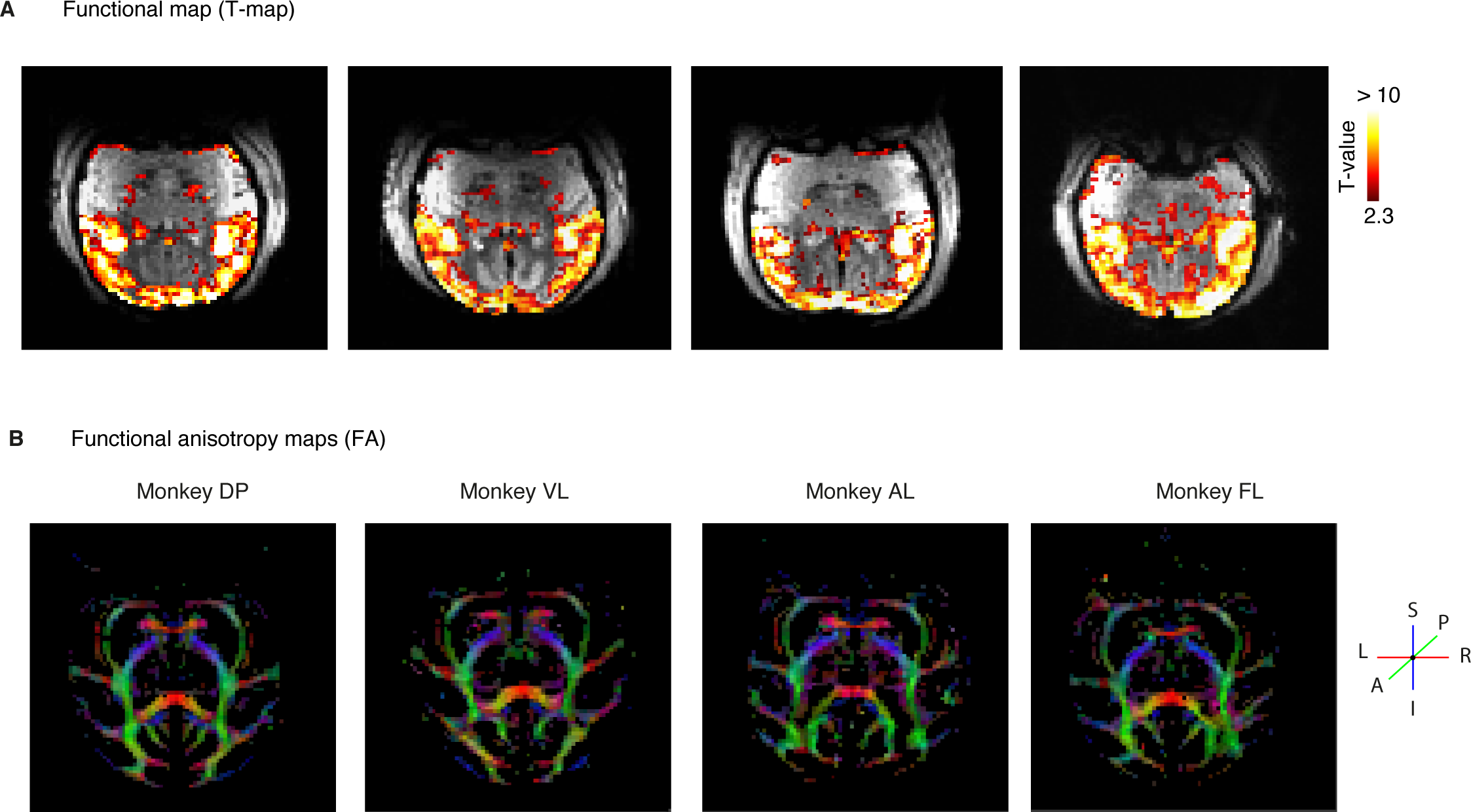
BOLD response and fractional anisotropy maps. **A**. Example echo-planar image shows the overall activation (T-value colormap range 2.3 < 10, FDR corr. q < 0.05) to the movie viewing of each subject. **B**. Example image from each NHP shows the directionally colored fractional anisotropy (FA).

**Supp. Fig. 2.**
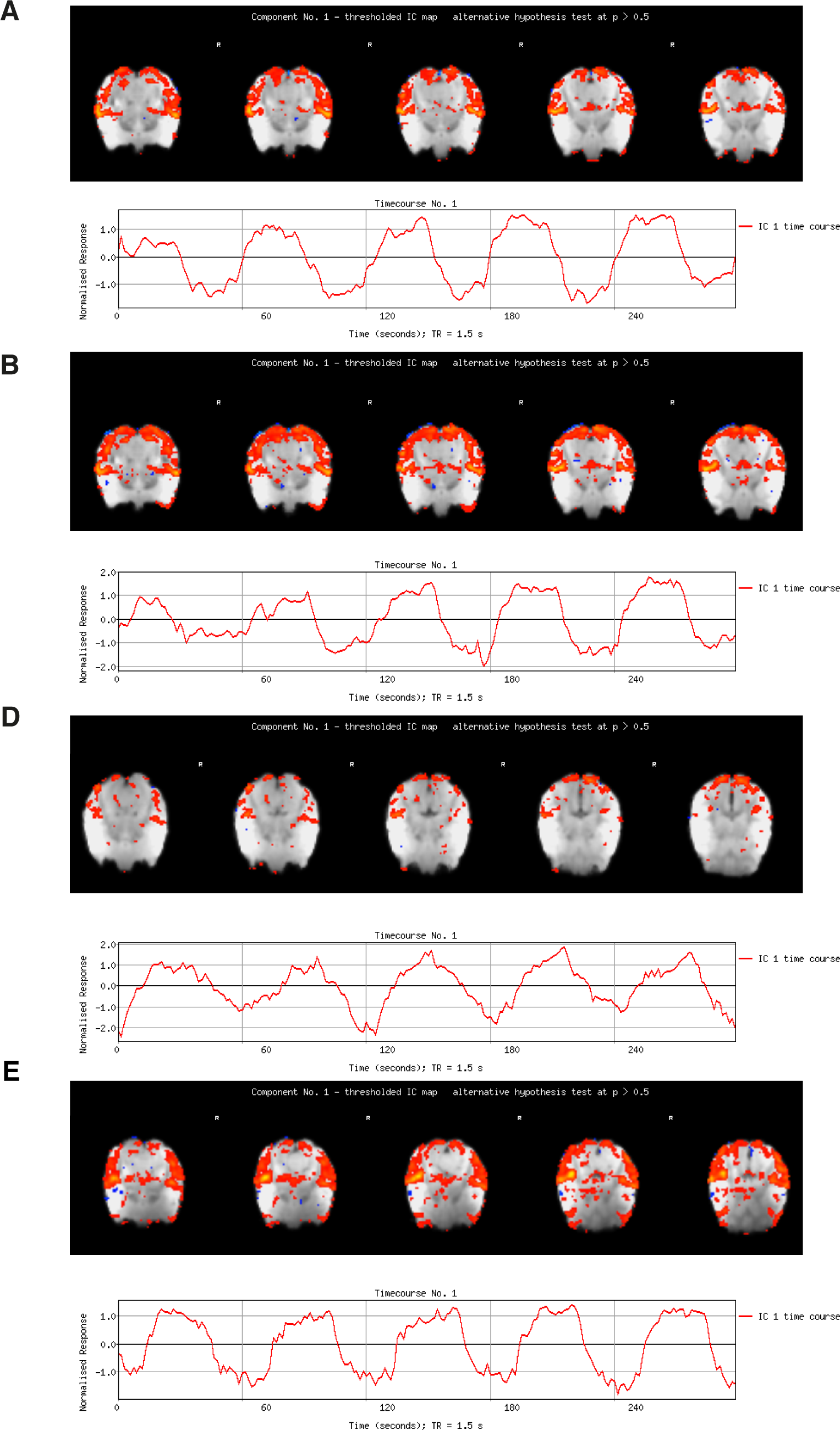
Independent component analyses and free-viewing pattern. **A**. Example slices of monkey AL showing the first ICA component pattern (10.33 % of explained variance, 5.68 % of total variance) of example run (5 mins duration). The independent component was observed on the first component of each ICA analyses. The plot below shows the independent component time course which generally reflects the stimulation rate of 30 secs ON and 30 secs OFF. **B**. Similar plots for monkey DP (9.41 % of explained variance, 5.25 % of total variance), **C** monkey FL (9.04 % of explained variance, 4.62 % of total variance) and **D** monkey VL (14.25 % of explained variance, 6.48 % of total variance). Each ICA map was threshold at a p-value of < 0.05.

**Supp. Fig. 3.**
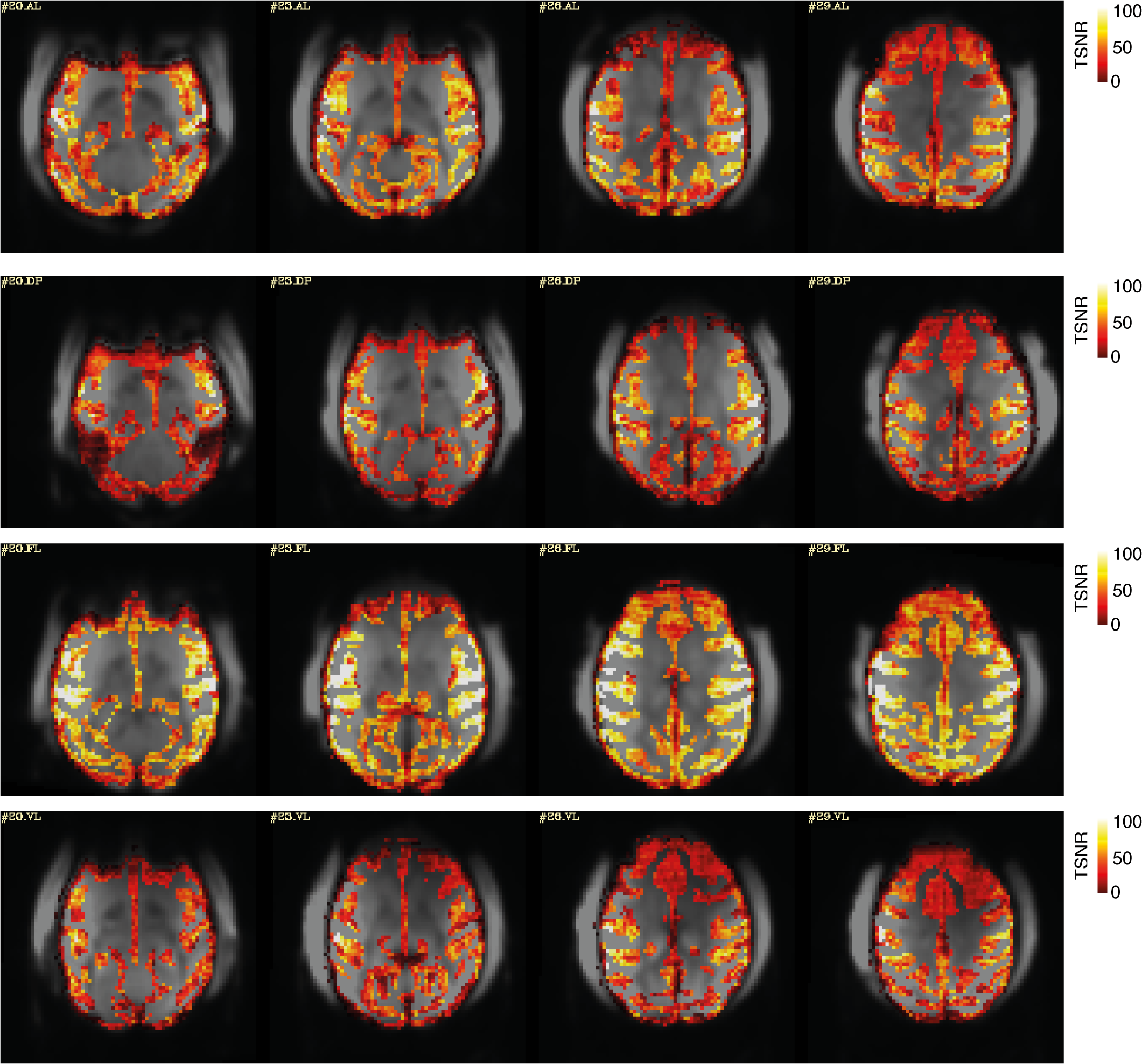
Temporal SNR maps of gray matter for each monkey. **A**. Four example slices of the temporal SNR maps for each monkey AL, DP, FL and VL. TSNR values clipped at 100 to ease visual inspection of maps across subjects.

**Supp. Fig. 4.**
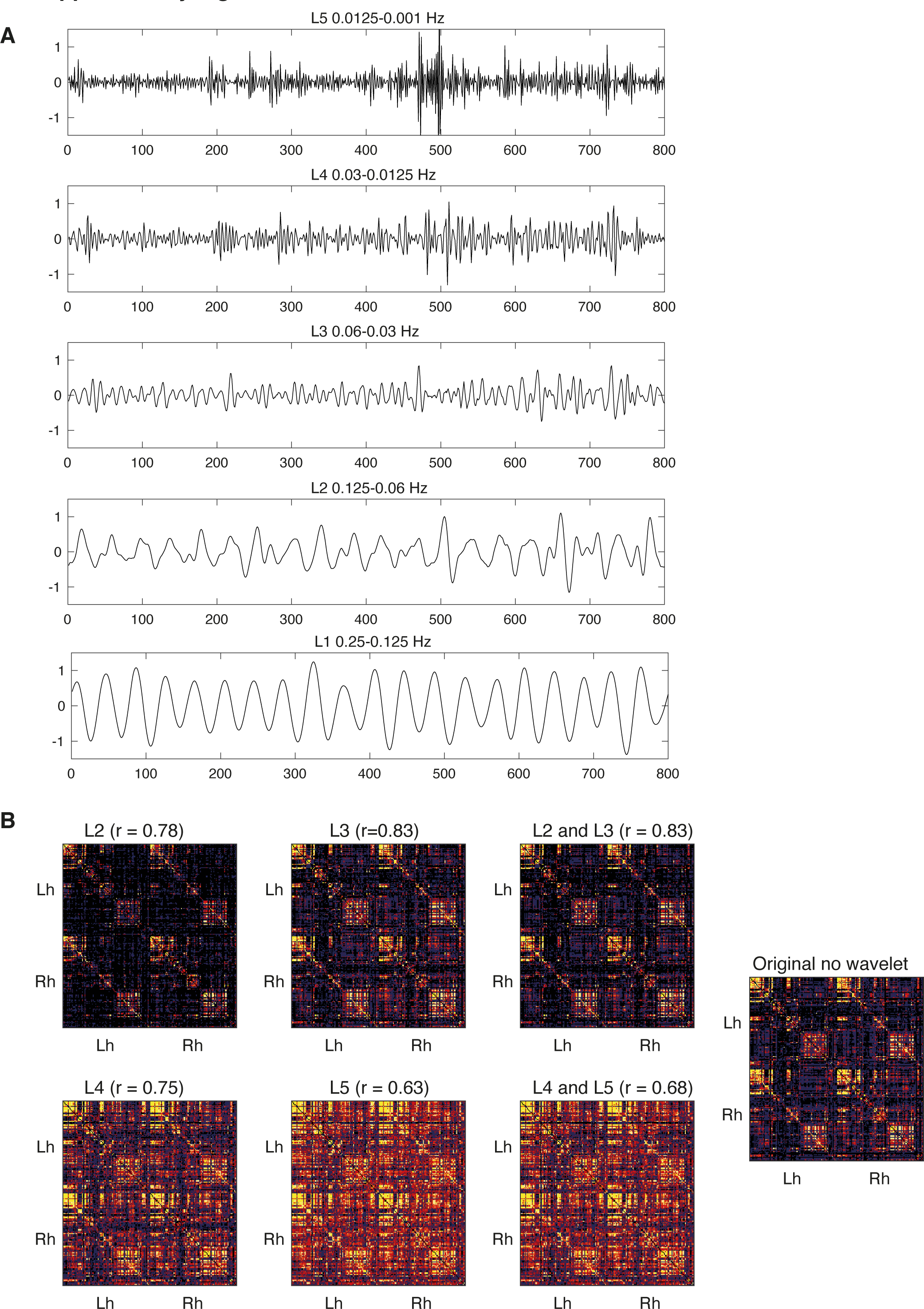
Wavelet decomposition of BOLD time series. **A**. Example time series with four runs concatenated using the maximal overlap discrete wavelet transform (MODWT) with the orthogonal Daubechies wavelet which resulted in 5 levels decompositions ranging from 0.25 to 0.001 Hz. For the construction of the free-viewing matrices, we concentrated on the relatively low-frequency range of 6 to 24 seconds, levels 2 (0.124 – 0.06) and level 3 (0.06 – 0.03). **B**. Constructed matrices of each wavelet Level 2 and Level 5 and their composites L2 and L3 and L4 and L5. The correlation coefficient (r) is between the level shown and the original matrix data without wavelet decomposition.

**Supp. Fig. 5.**
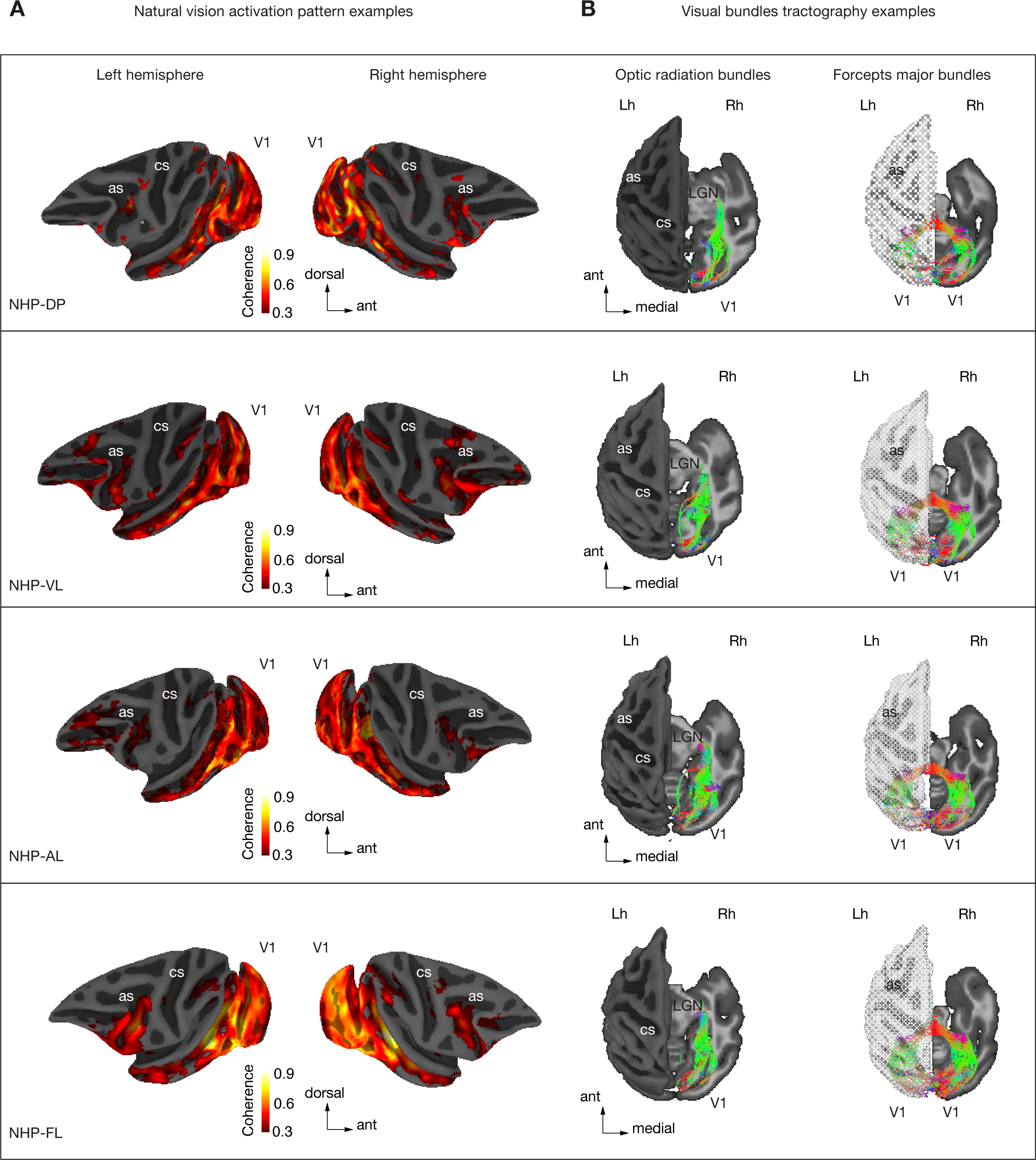
Functional mapping of visual activation during free-viewing and example visual bundle tracts. **A**. Lateral views of activation maps during natural free-viewing. Coherence maps are mapped into a semi-inflated brain surface of each subject. Regions with significant modulation (coherence > 0.35) included regions in ventrolateral lateral prefrontal cortex (vlPFC), frontal eye fields (FEF), lateral intraparietal area (LIP), visual regions (V1, V2, V3, V4), motion-sensitive regions (MT, MST, FST) and higher-level visual regions along the inferotemporal gyrus (TE, TEm, TPO, TEO, TEpd, IPa) among others. **B**. (Left) Example white-matter projections within the occipital cortex shown for each subject. Optic radiation bundle (OR) showing projections tracts from the LGN to visual cortex (V1). Forceps major (FM) bundle pathway originating in right V1 and projecting through the forceps major into left V1. For visualization, the plots showed a brain surface of the left hemisphere and were made semi-transparent.

**Supp. Fig. 6.**
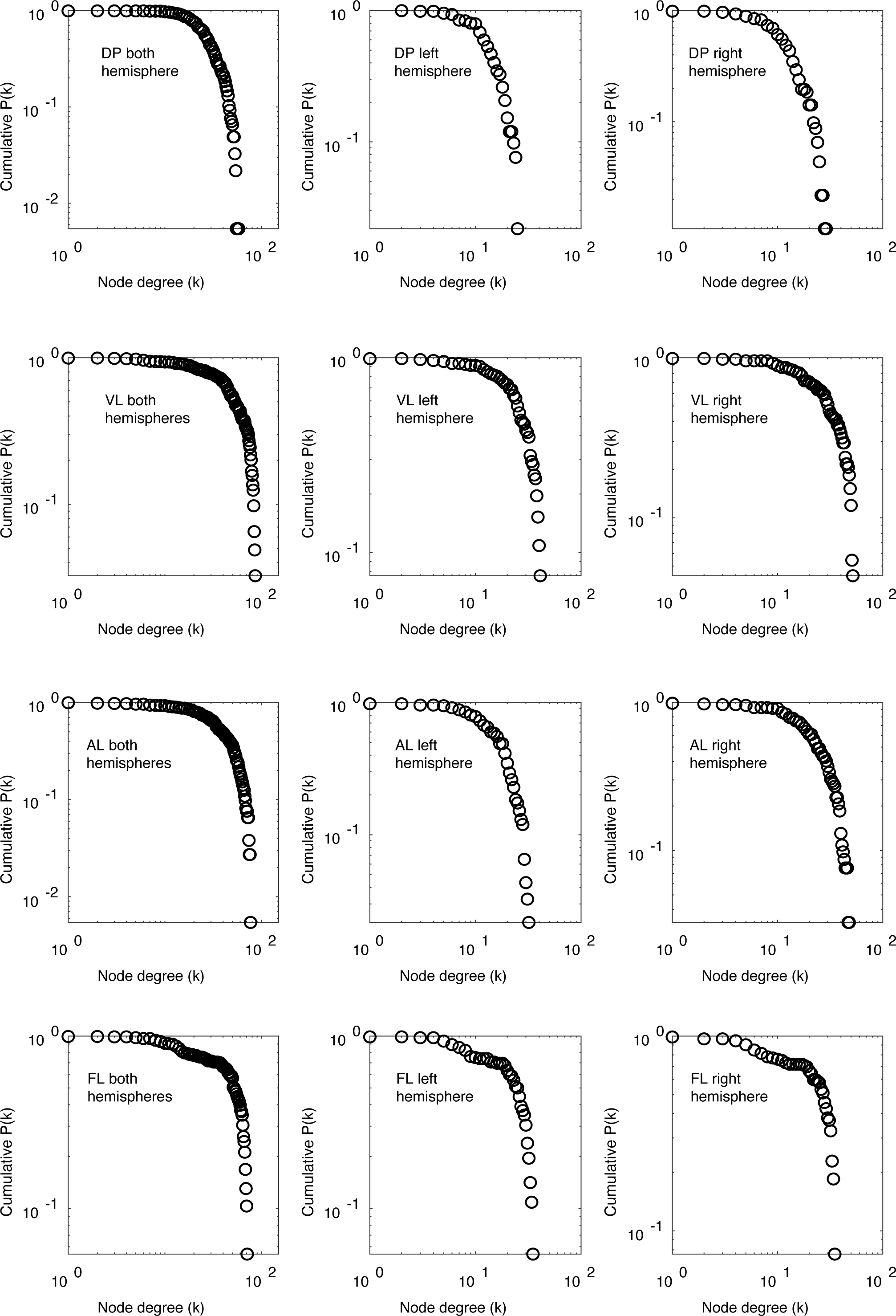
Free-viewing networks node degree distributions. Node degree distribution sample from a free-viewing network of each NHP and hemisphere.

**Supp. Fig. 7.**
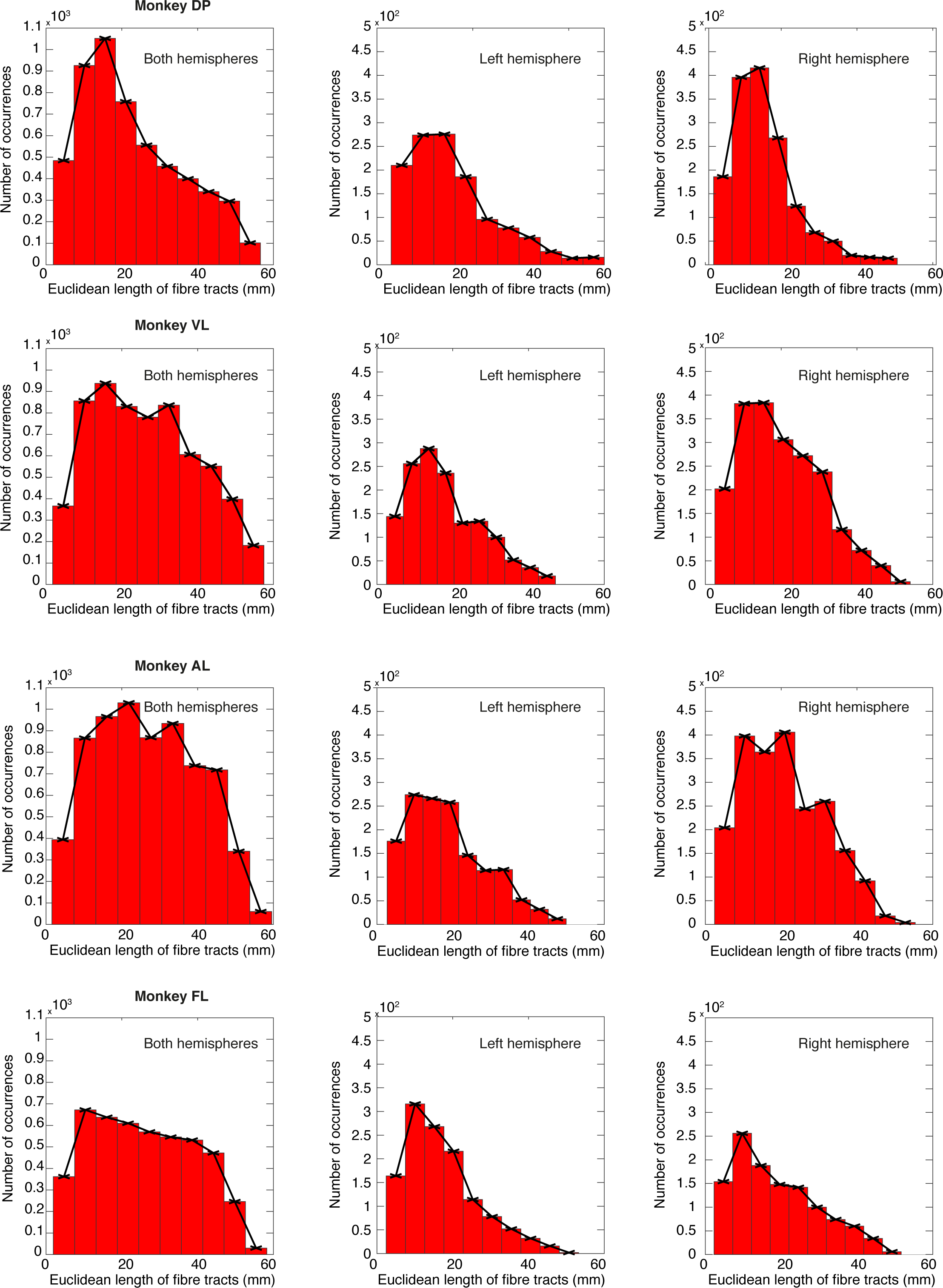
Free-viewing networks path length distributions. Path length distribution sample from a free-viewing network of each NHP and hemisphere.

**Supp. Fig. 8.**
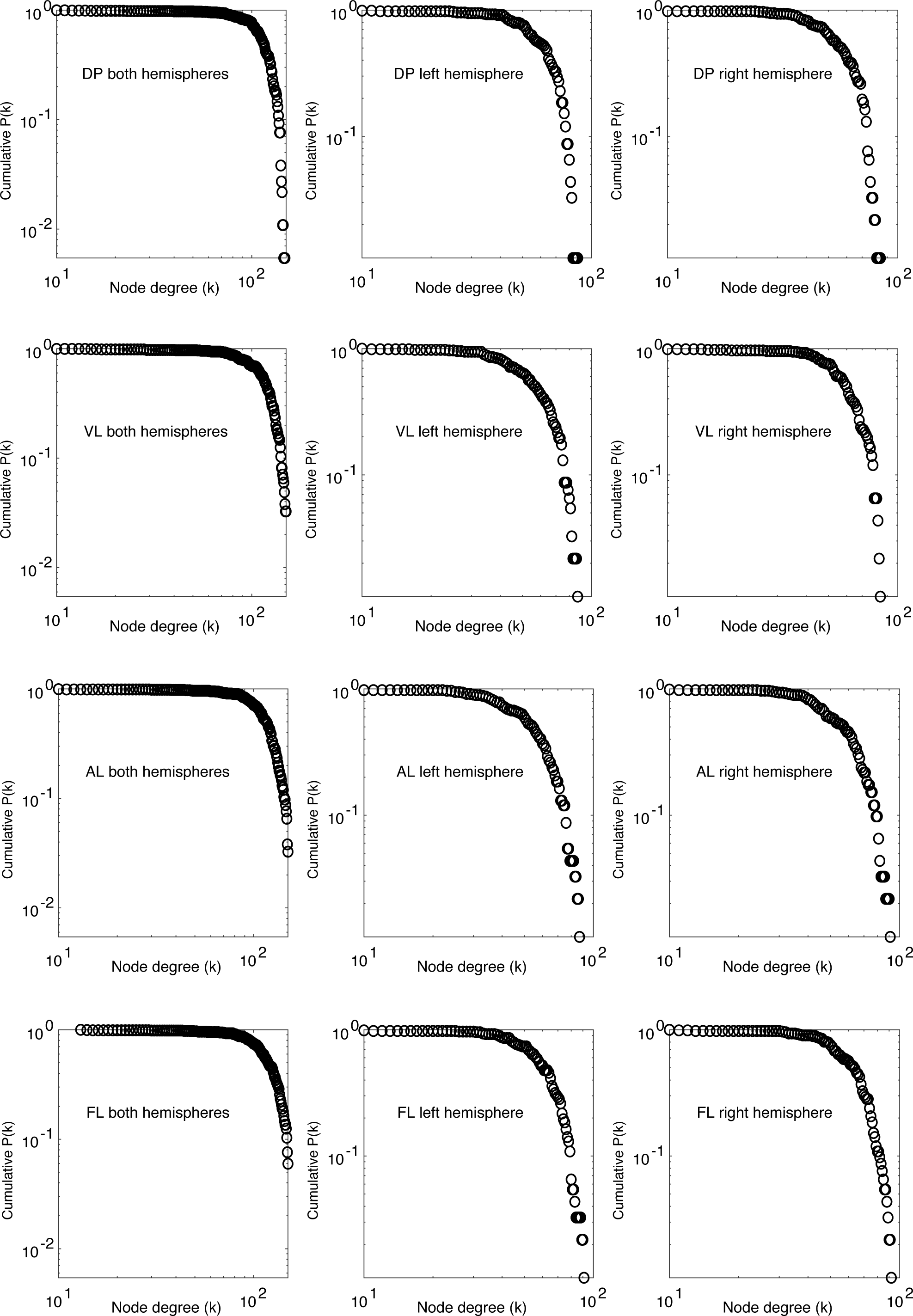
SN node degree distributions. Node degree distribution sample from structural networks of each NHP and hemisphere.

**Supp. Fig. 9.**
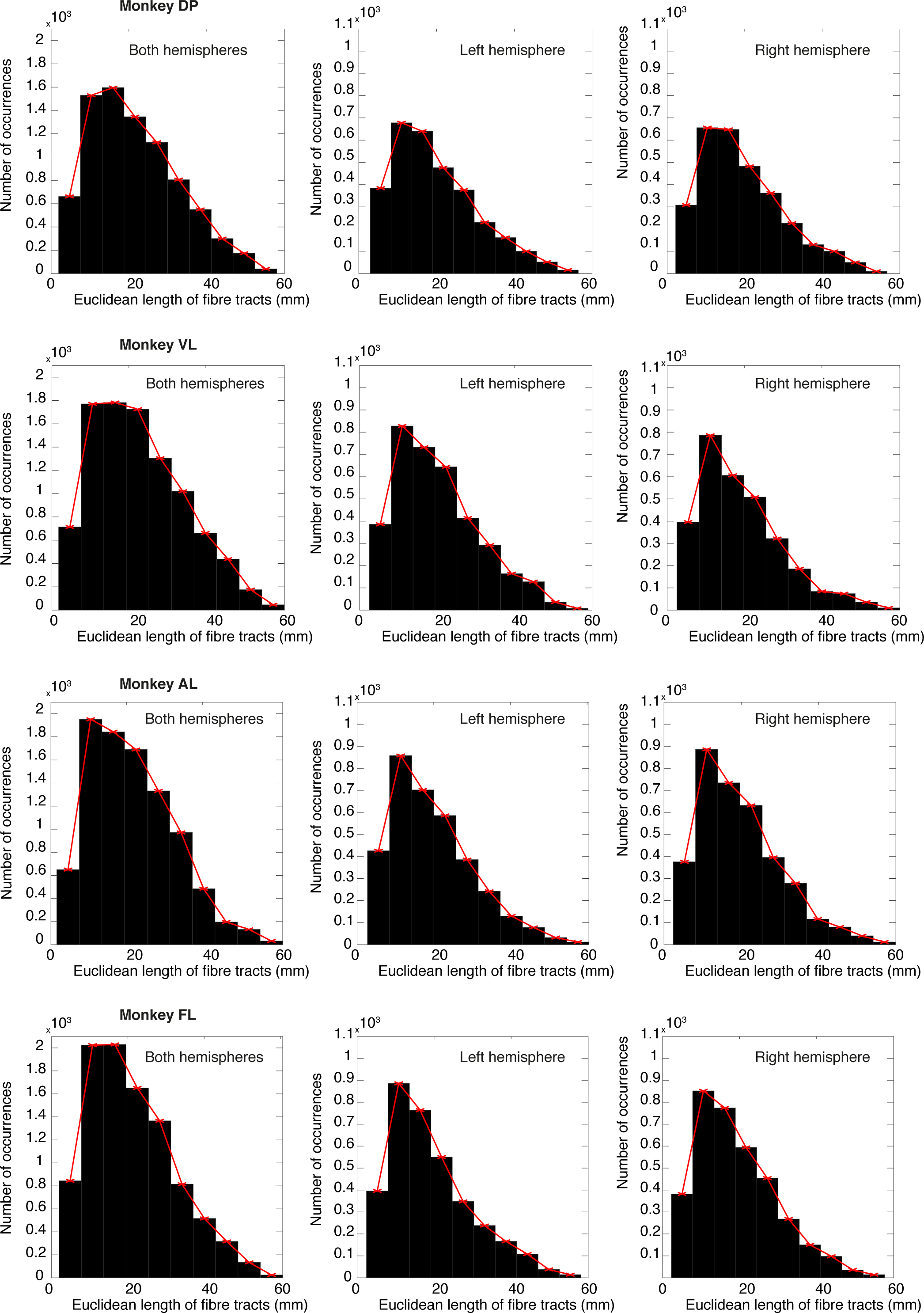
Structural path length distributions. Path length distribution sample from structural networks of each NHP and hemisphere.

**Supp. Fig. 10.**
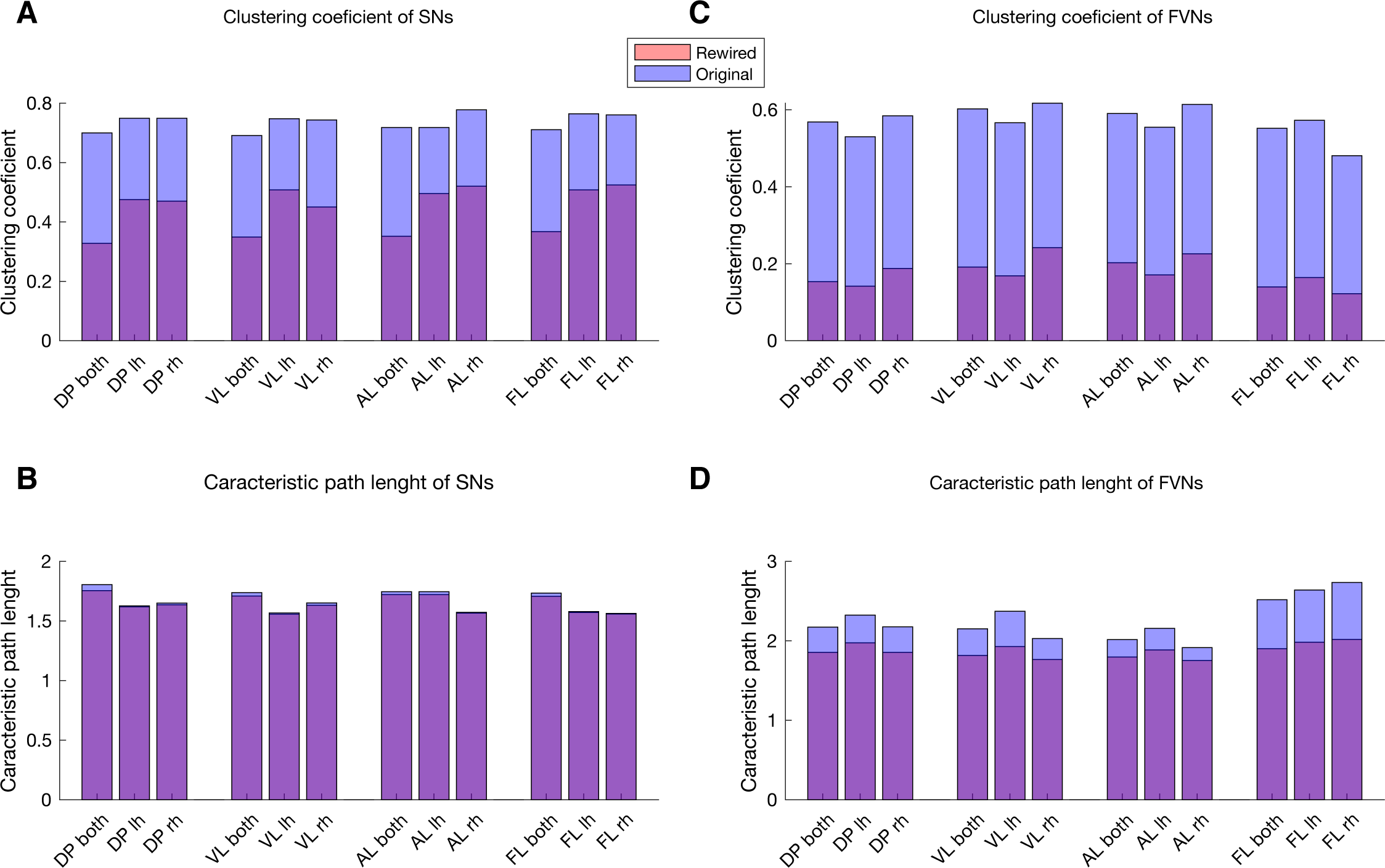
Clustering coefficient and characteristic path length. **A**. Bar plot of clustering coefficients of structural networks for both the original (blue) and rewired (red) of whole (both hemispheres), left and right alone. The transparency allows for the differential visualization between original and rewired networks. **B**. Same as in A for the characteristic path length of SN. **C**. Bar plot of clustering coefficients for Free-viewing networks **D**. Bar plot of characteristic path lengths for Free-viewing networks.

**Supp. Fig. 11.**
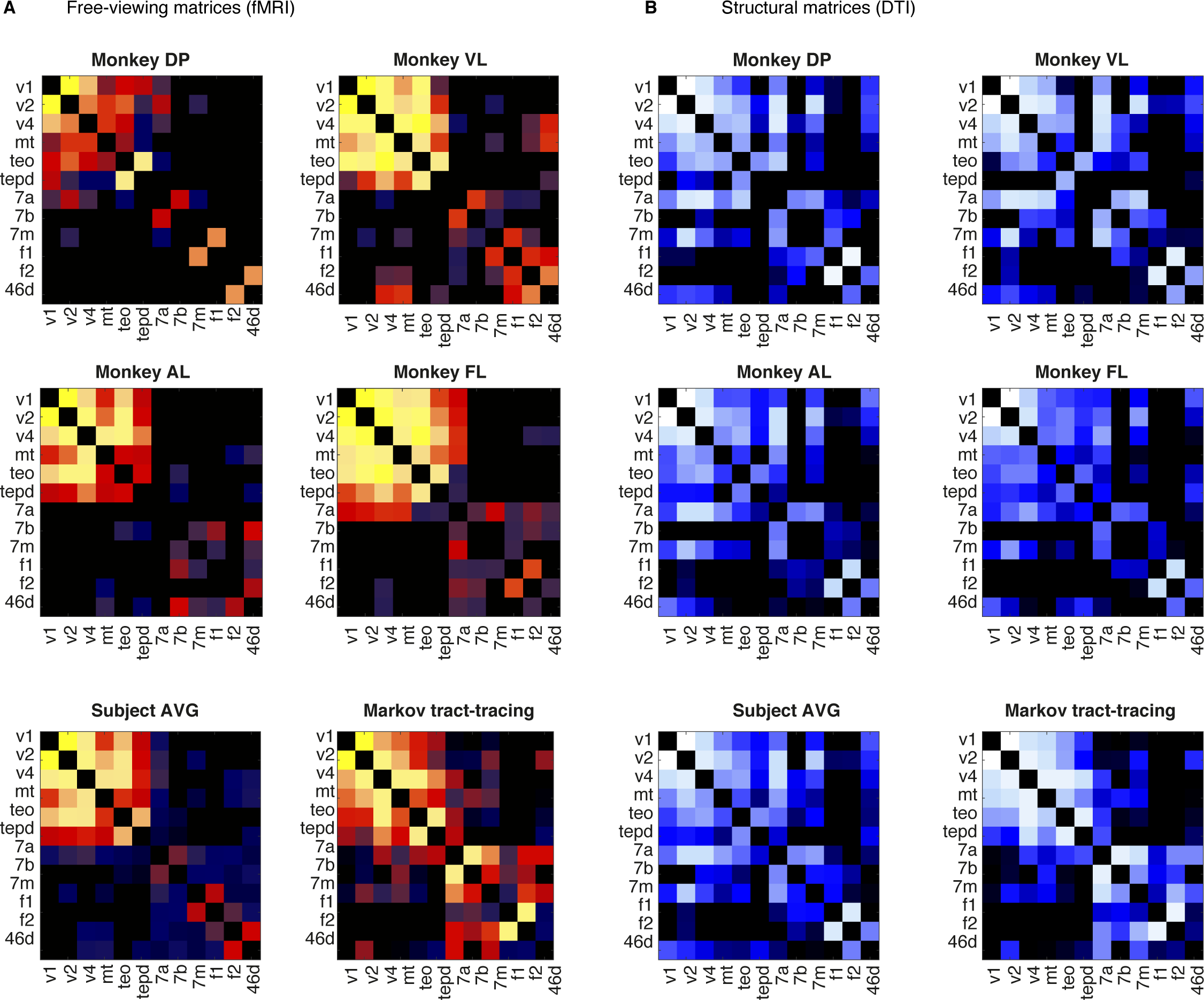
Structural and functional matrices organized according to overlapping regions from macaque tract-tracing matrix. A. DTI matrix for each NHP organized according to macaque tract-tracing data. B. fMRI-based matrices for each NHP organized according to tract-tracing data.

**Supp. Fig. 12.**
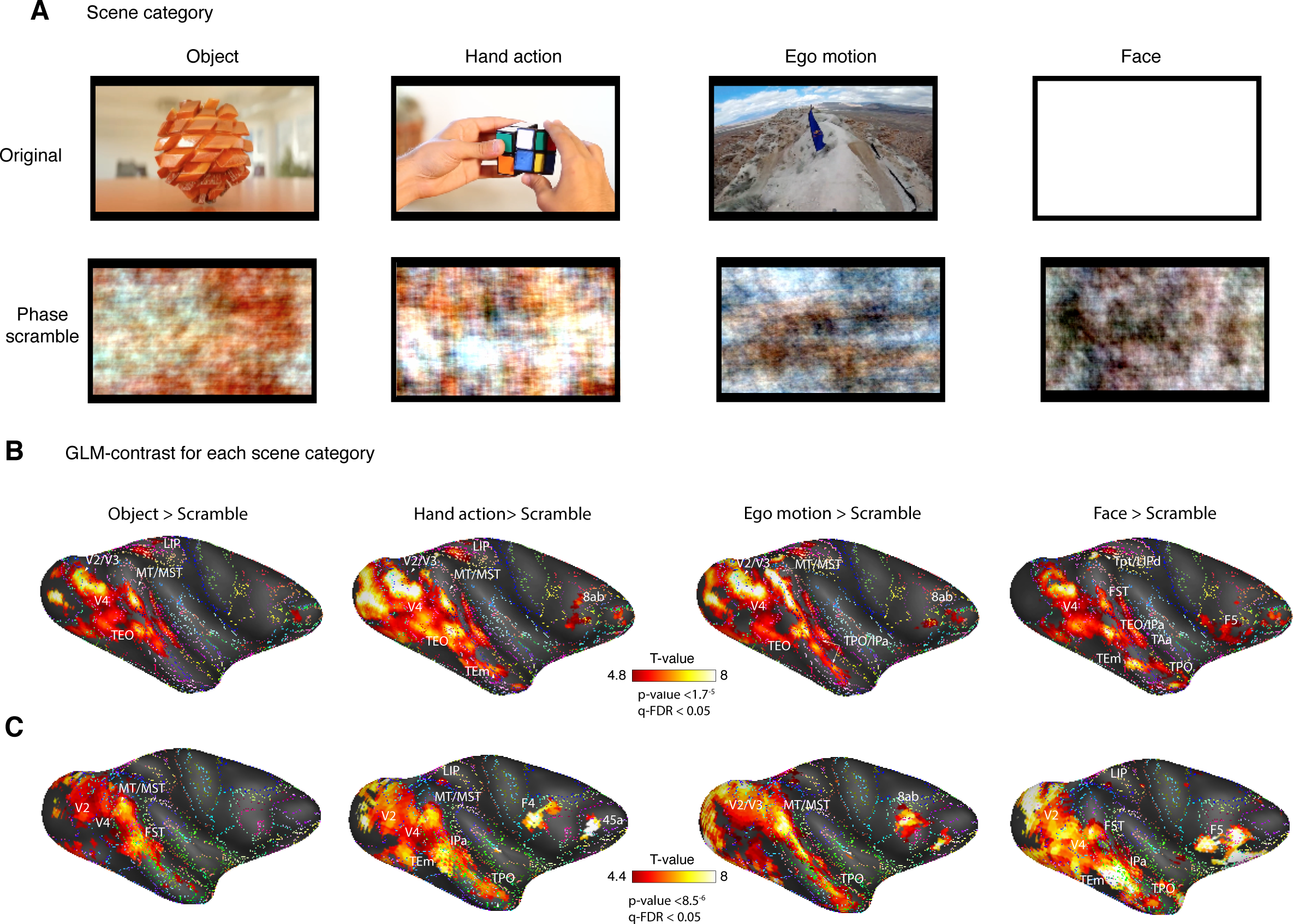
Movie segments stimuli used for contrast analyses between original scenes and phase scramble scenes. **A**. Example image frame of each movie clip: Ego-motion, Object, Face, Hand action. The movies were presented for 30 seconds followed by 15 secs of darkness. Each movie category contained four types of controls (Optic flow, phase scrambling, saliency contour and tile scrambling). **B**. GLM contrast (q-FDR < 0.05 corrected) between each scene category and correspondent phase scramble scenes shows activation patterns across the inferotemporal and frontal cortices for monkey DP. For each contrast, the t-value color range (4.8 > t < 8) allows the comparison across the activation magnitude of each scene category. **C**. Same contrast mapping for monkey AL.

**Supp. Fig. 13.**
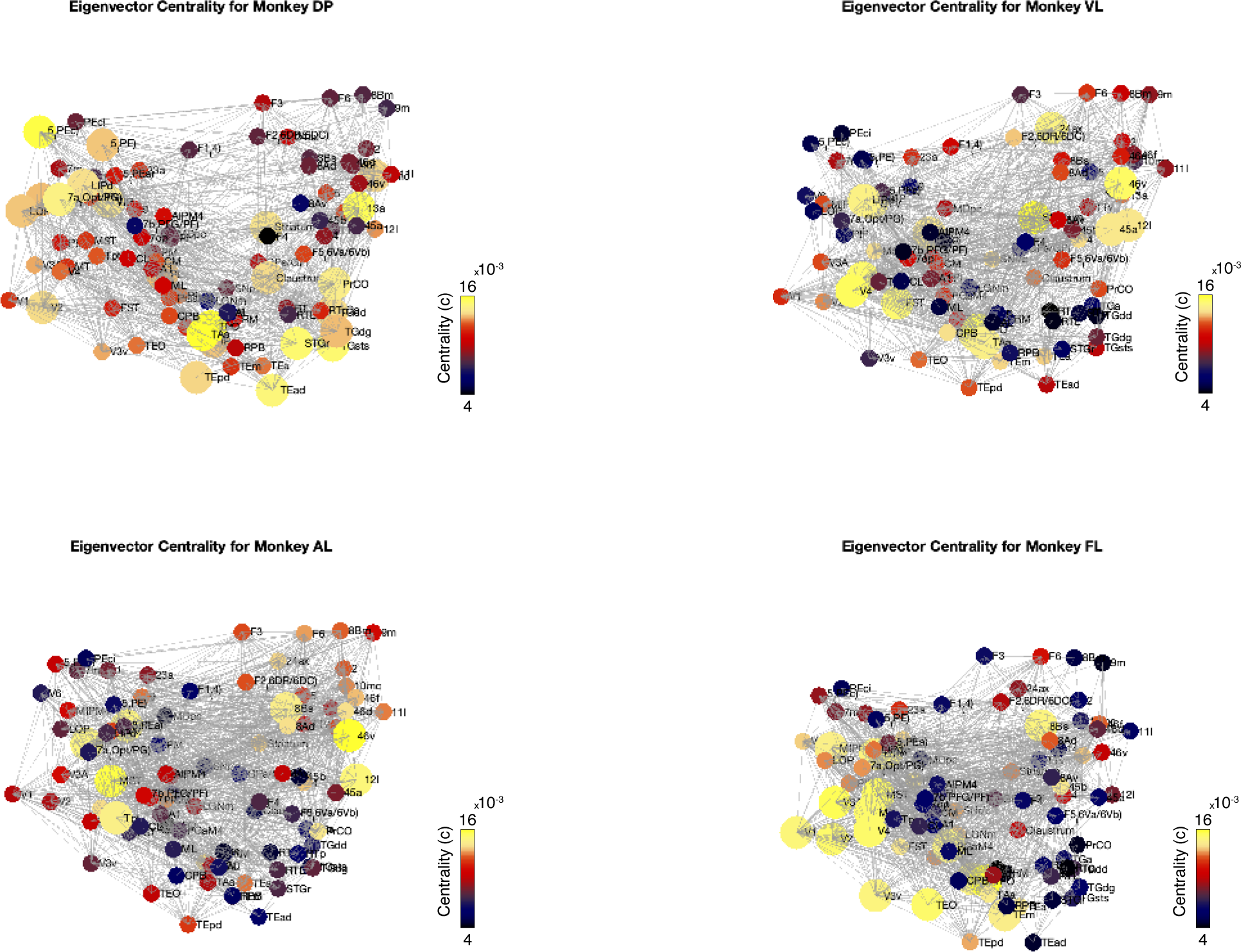
Network hubs for each monkey during free-viewing. Hubs with high eigenvector centrality highlighting regions with high-importance in the visual network during natural free-viewing. Hub regions for each monkey show eigenvector centrality (c) (c > 1x10^-4^; n highlight for size >= 1 Z-score) highlighting the central nodes during free-viewing.

**Supp. Table 1.** Small-world properties of structural brain networks obtained from binarized and rewired networks sharing the same degree distribution. Both, both hemispheres; lh, left hemisphere; rh, right hemisphere.

**Supp.Table2**
| subj.session.ru<br>n | component<br>num. | Explained<br>variance | Total<br>variance | Mean GM<br>TSNR |
| --- | --- | --- | --- | --- |
| V.GL1.R1 | 1 | 14.25% | 6.48% | 46.04 |
| V.GL1.R2 | 1 | 12.51% | 6.58% | 43.9 |
| V.GL1.R3 | 1 | 12.51% | 6.33% | 46.83 |
| DP.GI2.R1 | 1 | 9.41% | 5.25% | 46.36 |
| DP.GI2.R2 | 1 | 12.27% | 6.29% | 48.71 |
| DP.GI2.R3 | 1 | 8.40% | 5.04% | 51.37 |
| DP.GI2.R4 | 1 | 7.22%% | 4.24% | 44.69 |
| F.MB1.R1 | 1 | 9.05% | 4.62% | 59.62 |
| A.PR1.R1 | 1 | 10.33% | 5.68% | 81.99 |
| A.PR1.R2 | 1 | 9.38% | 5.24% | 57.92 |

**Supp. Table 2.** Network properties. Small-world properties of structural brain networks obtained from binarized and rewired networks sharing the same degree distribution. Both, both hemispheres; lh, left hemisphere; rh, right hemisphere.

| ***NHP-Hem-Net*** | ***Edge***<br>***densit***<br>***y*** | ***Clustering***<br>***coefficient***<br>***rewired*** | ***Clustering***<br>***coefficient*** | ***Characteristic***<br>***path length,***<br>***rewired*** | ***Characteristic***<br>***path length*** |
| --- | --- | --- | --- | --- | --- |
| DP-both-SN | 0.2469 | 0.3283 | 0.7003 | 1.7550 | 1.8052 |
| DP-lh-SN | 0.3829 | 0.4757 | 0.7492 | 1.6190 | 1.6274 |
| DP-rh-SN | 0.3660 | 0.4705 | 0.7494 | 1.6362 | 1.6503 |
| VL-both-SN | 0.2916 | 0.3496 | 0.6911 | 1.7092 | 1.7385 |
| VL-lh-SN | 0.4448 | 0.5084 | 0.7477 | 1.5573 | 1.5676 |
| VL-rh-SN | 0.3703 | 0.4504 | 0.7436 | 1.6321 | 1.6522 |
| AL-both-SN | 0.2812 | 0.3520 | 0.7185 | 1.7218 | 1.7461 |
| AL-lh-SN | 0.4233 | 0.4957 | 0.7185 | 1.7218 | 1.7461 |
| AL-rh-SN | 0.4350 | 0.5205 | 0.7777 | 1.5670 | 1.5730 |
| FL-both-SN | 0.2941 | 0.3674 | 0.7112 | 1.7069 | 1.7342 |
| FL-lh-SN | 0.4300 | 0.5081 | 0.7645 | 1.5714 | 1.5789 |
| FL-rh-SN | 0.4436 | 0.5250 | 0.7606 | 1.5584 | 1.5636 |
| DP-both-FN | 0.1595 | 0.1534 | 0.5683 | 1.8550 | 2.1729 |
| DP-lh-FN | 0.1476 | 0.1419 | 0.5297 | 1.9763 | 2.3239 |
| DP-rh-FN | 0.1861 | 0.1877 | 0.5844 | 1.8564 | 2.1770 |
| VL-both-FN | 0.1884 | 0.1913 | 0.6023 | 1.8164 | 2.1519 |
| VL-lh-FN | 0.1665 | 0.1686 | 0.5663 | 1.9288 | 2.3725 |
| VL-rh-FN | 0.2410 | 0.2418 | 0.6169 | 1.7659 | 2.0293 |
| AL-both-FN | 0.2053 | 0.2028 | 0.5904 | 1.7978 | 2.0170 |
| AL-lh-FN | 0.1727 | 0.1710 | 0.5547 | 1.8860 | 2.1571 |
| AL-rh-FN | 0.2563 | 0.2258 | 0.6137 | 1.7518 | 1.9160 |
| FL-both-FN | 0.1389 | 0.1397 | 0.5519 | 1.9029 | 2.5182 |
| FL-lh-FN | 0.1503 | 0.1644 | 0.5727 | 1.9833 | 2.6407 |
| FL-rh-FN | 0.1388 | 0.1224 | 0.4806 | 2.0177 | 2.7350 |

